# Paired Tumor Biopsies Reveal Spatiotemporal Myeloid Remodeling After Local Chemotherapy in Glioblastoma

**DOI:** 10.64898/2026.05.13.725006

**Authors:** Anthony J. Tang, Matthew R. Warren, Peter J. Chabot, Damian E. Teasley, Nicholas B. Dadario, Angeliki Mela, Misha Amini, Nathaniel W. Rolfe, Michael G. Argenziano, Colin P. Sperring, Alan X. Chen, Nkechime Ifediora, Ashwin Viswanathan, Melanie Kristt, Athanassios Dovas, Brianna Pereira, Abby Brand, August Kahle, Nelson Humala, Clara Stucke, Julia Furnari, Corina Kotidis, Wenting Zhao, Zhouzerui Liu, Shikun Wang, Hannah Haile, Nadine Khoury, Arjun R. Adapa, Nathan J. Winans, Nina Yoh, Justin A. Neira, Kristin R. Swanson, Brian J. A. Gill, Peter Sims, Osama Al Dalahmah, Jack Grinband, Liang Lei, Peter Canoll, Jeffrey N. Bruce

## Abstract

**Background:** Standard-of-care chemotherapy for glioblastoma induces inflammation that may contribute to disease recurrence. Although recurrent tumors are enriched with myeloid cells, the early cellular response to chemotherapy and the mechanisms that initiate this inflammatory remodeling remain poorly understood. In particular, it is unknown how neoplastic and tumor-associated myeloid populations respond during the immediate post-treatment period, and whether tumor-associated myeloid cells directly experience chemotherapy-induced genotoxic stress that contributes to inflammatory state transitions.

**Methods:** We performed sequencing-based analysis of neoplastic and immune populations following topotecan exposure using multiple complementary model systems and time points. We first analyzed MRI-localized, paired pre- and post-treatment biopsies from a first-in-human trial of 28-day convection enhanced delivery (CED) of topotecan (n=5) using cell-type-deconvolved bulk-RNA-sequencing and immunofluorescence. We then treated syngeneic murine gliomas using an *in vivo* model of CED-topotecan and measured acute 3-day and 7-day treatment responses by single cell RNA-sequencing. We additionally conducted sequencing analysis of patient-derived slice cultures and *in vitro* human microglial and glioma cell lines following 24-hour topotecan treatment.

**Results:** In paired human biopsies, CED-topotecan induced spatially restricted transcriptional remodeling within the infusion zone, characterized by suppression of proliferative tumor programs and enrichment of inflammatory, interferon, hypoxia, and mesenchymal signatures. Cell-type deconvolution and immunofluorescence linked this response to myeloid remodeling, including enrichment of monocyte-derived tumor-associated macrophage states, increased MARCO-positive myeloid populations, and pH2AX-positive genotoxic stress within Iba1-positive myeloid cells. In the murine CED model, topotecan prolonged survival and reduced tumor cellularity, while also inducing inflammatory and DNA-damage programs in tumor-associated macrophages that evolved by 7-days toward hypoxia, angiogenesis, TGF-β signaling, and mesenchymal/tissue-remodeling programs. Human slice culture and *in vitro* microglial systems confirmed stress-coupled inflammatory and DNA-damage responses in human myeloid cells.

**Conclusions:** Chemotherapy exposure induces a spatially structured inflammatory myeloid response characterized by early genotoxic stress and inflammatory activation, with later emergence of mesenchymal and tissue-remodeling macrophage programs. Across model systems, our analysis supports a model in which chemotherapy-associated damage in both tumor and myeloid cells contributes to an evolving inflammatory microenvironment after treatment.

## INTRODUCTION

Glioblastoma (GBM) recurrence after cytotoxic therapy is frequently accompanied by inflammatory and mesenchymal remodeling of the tumor microenvironment (1–3). Tumor-associated myeloid cells, including resident microglia and infiltrating monocyte-derived macrophages, are central regulators of this process, shaping inflammatory signaling, immune suppression, mesenchymal transition, and therapeutic resistance (2, 4–11). Mesenchymal-like GBM states are particularly enriched for inflammatory myeloid programs and are associated with disease progression and poor clinical outcomes (2, 3, 12–15). Although chemotherapy has traditionally been interpreted primarily through its effects on tumor cells, including reduced proliferation, DNA damage, and cell death, cytotoxic therapy also exposes surrounding immune and stromal compartments to treatment-associated stress (3, 16).

A recent phase Ib trial in recurrent GBM evaluating chronic convection-enhanced delivery (CED) of topotecan provided important insight into post-treatment myeloid dynamics (17). Histologic analyses of paired pre- and post-treatment clinical specimens showed a significant decrease in proliferative tumor markers after treatment, including reduced Ki67+ and SOX2+ tumor cells (17). These specimens also revealed increased inflammatory features within the tumor microenvironment, including enrichment of inflammatory and mesenchymal transcriptional signatures and evidence of activated CD68+ myeloid cells on immunohistochemical analysis (17), indicating that local chemotherapy exerts parallel cytotoxic and microenvironmental effects. However, the temporal evolution of this response, the specific cell populations involved, and the molecular programs that drive therapy-associated inflammatory remodeling of the tumor microenvironment remain poorly understood.

Previous studies of treatment response and recurrence biology in GBM have relied on specimens obtained at recurrence, often compared with pretreatment primary tumors, archival specimens, or unmatched controls (18–20). Although these approaches have provided important insights into the biology of recurrent disease, specimens obtained at recurrence reflect the cumulative effects of treatment, tumor regrowth, and ongoing tumor evolution, making it difficult to define the primary cellular and molecular responses elicited by chemotherapy (18, 21, 22). As a result, the immediate post-treatment interval--when chemotherapy-induced stress, inflammation, and microenvironmental remodeling may first become evident--remains poorly characterized. Paired pre- and immediate post-treatment biopsies from the same patient offer a more direct view of how tumor and immune compartments respond to therapy before clinical or radiographic recurrence occurs. Indeed, serial tissue sampling has been advocated as necessary for GBM drug development because conventional radiographic and survival endpoints provide limited insight into tissue-level drug effects and mechanisms of treatment failure (23–28). The paired, MRI-localized clinical specimens from the CED-topotecan trial therefore provide a rare opportunity to interrogate the immediate effects of local chemotherapy in human GBM and to characterize early inflammatory and microenvironmental changes that may precede recurrence.

In this study, we used paired human biopsies and complementary experimental models to characterize the temporal and cellular architecture of the myeloid response to topotecan exposure in GBM. We hypothesized that chemotherapy-associated inflammation reflects a staged myeloid remodeling process, with early genotoxic stress and inflammatory activation followed by later enrichment of macrophage-associated mesenchymal and tissue-remodeling programs. To map this post-treatment myeloid landscape, we used CED-topotecan as a core model system, because local delivery enables interrogation of direct chemotherapy exposure in a clinically relevant setting while minimizing confounds from systemic drug effects. We performed spatiotemporal differential expression analysis and cell-type compositional deconvolution of MRI-localized bulk RNA-seq biopsies from 5 patients with recurrent GBM treated with CED-topotecan for 28 days (17) to define the spatially localized transcriptional and compositional changes associated with drug exposure. Because these clinical samples captured the response after 28 days of exposure, we used syngeneic glioma mice treated with CED-topotecan for 3 and 7 days to resolve earlier myeloid responses *in vivo*. We further used *ex vivo* human GBM slice cultures and isolated human microglial models to interrogate acute responses in human tissue and determine the direct effects of topotecan on microglia. Together, our findings support a model in which local chemotherapy reshapes the GBM tumor-immune microenvironment through effects on both neoplastic and myeloid populations, linking genotoxic stress to spatially and temporally organized inflammatory remodeling after treatment.

## RESULTS

### 1. CED-topotecan induces spatially localized mesenchymal and inflammatory remodeling in human recurrent GBM

We first asked whether chronic CED-topotecan is associated with systems-level remodeling of the recurrent GBM transcriptome, and whether such remodeling is spatially restricted to the maximal drug infusion zone. We analyzed bulk RNA-seq data from matched pre- and post-treatment biopsies collected at catheter implantation and explantation, respectively, and stratified post-CED specimens by whether they were obtained within the maximal infusion zone (post-CED IN) or outside of it (post-CED OUT) (**Figure 1A, Figure S1A**) (17). We projected gene-wise log-2-fold-change values for post-CED IN and post-CED OUT samples, each calculated relative to pre-CED, in the same coordinate space. As shown in **Figure 1B**, many genes changed in a similar direction and magnitude across spatial compartments (top right and bottom left quadrants, representing genes up- or down-regulated in both zones, respectively; only genes with FDR-adjusted *p* < 0.05 shown), whereas a substantial subset showed discordant differential expression inside versus outside the infusion zone (top left and bottom right quadrants). CED-topotecan induced region-specific transcriptional changes related to direct drug exposure. Hallmark gene set enrichment analysis using the MSigDB pathways (**Supp. Figure S1B**) showed that both post-CED IN and post-CED OUT were enriched for inflammatory and cytokine signaling programs, including IL-6/JAK/STAT3, TNF-α signaling via NF-κB, apoptosis, hypoxia, and other stress-associated pathways (ROS, Epithelial/Mesenchymal Transition), with negative enrichment for proliferative and metabolic programs (mitotic spindle, KRAS, peroxisome). The infusion zone-specific transcriptional signature (top left and bottom right quadrants) showed a distinct inflammatory and anti-proliferative profile, characterized by high interferon alpha/gamma response and allograft rejection with low proliferative pathway scores (G2-M checkpoint, E2F targets).

**Figure 1:**
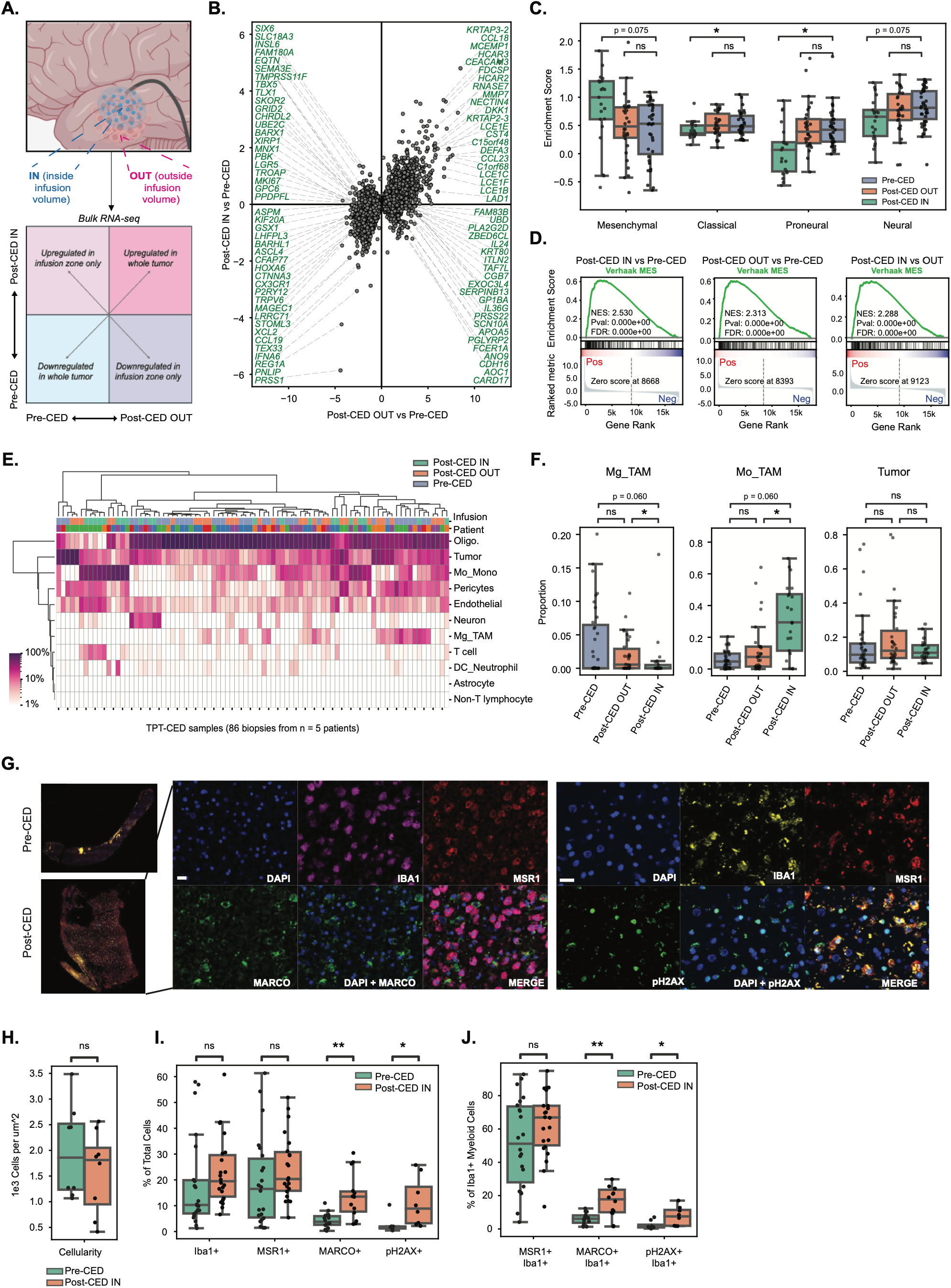
Spatially resolved transcriptomic remodeling of the recurrent GBM microenvironment following CED-topotecan. **(A)** Study design and spatial biopsy acquisition from patients with recurrent GBM treated with CED of topotecan, including pre-CED biopsies and post-CED biopsies stratified by location inside or outside the maximal MRI-defined infusion zone. **(B)** Differential gene-expression analysis comparing pre-CED, post-CED-IN, and post-CED-OUT biopsies, highlighting induction of immune-associated chemokines and suppression of proliferative tumor programs within the infusion zone. **(C)** Signature scoring of canonical Verhaak subtypes across patient biopsies demonstrating depletion of proneural and classical programs within post-CED-IN tissue. **(D)** Gene set enrichment analysis scores for the Verhaak mesenchymal signature comparing post-CED-IN versus pre-CED, post-CED-OUT versus pre-CED, and post-CED-IN versus post-CED-OUT tissue. **(E)** Digital cytometry–based inference of cell-type composition across pre-CED, post-CED-IN, and post-CED-OUT biopsies. **(F)** Biopsy-level statistical comparisons of compositional profiles depicting increased Mo-TAMs and reduced Mg-TAMs within the infusion zone. **(G)** Representative images of multiplex immunofluorescence staining for DAPI, IBA1, MSR1, pH2AX, and MARCO in paired pre-CED and post-CED-IN biopsies. Scale bar = 20 µm. **(H–J)** Quantification of overall cellularity (H), total marker abundance (I), and phenotypic distribution of IBA1+ myeloid cells (J) following treatment. Statistics were calculated using linear models with patient as fixed effects, adjusting for patient-specific differences across multiple biopsies from 5 patients. P values were adjusted within each contrast using the Benjamini–Hochberg false discovery rate procedure. *q < 0.05; **q < 0.01.

We also scored each biopsy for canonical bulk GBM subtype signatures (mesenchymal, classical, proneural, and neural) defined by Verhaak et al. (30). Sample-level signature scoring showed decreased proneural and classical signatures in post-CED IN biopsies relative to pre-CED biopsies (patient-adjusted *p* = 0.0099 and 0.015, respectively) **(Figure 1C)**. Differential expression analysis with patient-level correction followed by GSEA confirmed enrichment of the Verhaak mesenchymal signature in post-CED IN tissue relative to pre-CED and post-CED OUT tissue **(Figure 1D)** (FDR-adjusted *p* < 0.001 for all comparisons). These complementary analyses support a spatially restricted subtype shift, with post-CED OUT samples showing attenuated changes relative to infusion-zone biopsies. Together, these findings indicate that local topoisomerase inhibition induces spatially restricted inflammatory and proneural/classical to mesenchymal remodeling within drug-exposed tumor regions.

### 2. Bulk deconvolution and immunofluorescence demonstrate mesenchymal monocyte-derived TAM enrichment and broad DNA damage within the infusion zone

While bulk RNA-seq analysis captures treatment-associated changes in the tumor microenvironment transcriptome, these profiles reflect average expression across all cells within a sample and may be influenced by differences in cellular composition between samples. We therefore assessed compositional changes induced by CED-topotecan and compared compositional profiles across infusion zone regions using CIBERSORTx bulk deconvolution **(Figure 1E-F)** (29). Unsupervised clustering of inferred cell-type fractions showed that biopsies segregated primarily by treatment group (pre-CED, post-CED IN, post-CED OUT) rather than by patient, indicating a shared treatment-associated compositional shift across individuals (**Figure 1E**). We focused subsequent statistical comparisons on tumor cells and the major myeloid compartments, including monocyte-derived tumor-associated myeloid cells (Mo-TAMs), which have been linked to chronic inflammation and mesenchymal GBM states (14), and microglia-like tumor-associated myeloid cells (Mg-TAMs), which are associated with homeostatic and acute inflammatory states. Tumor-cell signal remained relatively stable across biopsy regions, whereas Mo-TAMs were enriched and Mg-TAMs were depleted following treatment within the infusion zone (post-CED IN) compared to outside (post-CED OUT) (**Figure 1F**; patient-adjusted *p =* 0.031 for both comparisons). These myeloid shifts were most pronounced within the infusion zone, consistent with the inflammatory enrichment at the bulk-transcriptional level and suggesting a major contribution from treatment-induced myeloid reorganization. To support these inferred compositional changes, we compared CIBERSORTx-derived cell-type abundance estimates with multiplex immunofluorescence–based quantification of matched biopsy tissue. These analyses showed concordance between transcriptomic deconvolution and orthogonal tissue-level staining measurements, supporting the validity of the inferred myeloid remodeling patterns **(Supp. Figure S2C)**.

To further delineate cell type-specific contributions to the bulk transcriptional signal, we employed the CIBERSORTx High-Resolution algorithm to infer cell type-specific expression profiles for each biopsy. We inspected biopsy-wise expression profiles for tumor, Mo-TAM, and Mg-TAM cell populations, thereby approximating cell type-specific expression profiles from the bulk data **(Supp. Figure S1F)**. Post-CED IN tumor signatures were enriched for hypoxia-adaptive and mesenchymal-associated genes, including *HK2*, *LPL*, *HILPDA*, *EGLN3*, *NDRG1*, *PRRX1*, and *TNC* (**Supp. Figure S1C**). Mo-TAM signatures in both post-CED IN and post-CED OUT samples were marked by expression of *MARCO*, *CLEC5A*, *TREM1*, *MMP19*, *PTGS2*, and *ADM* **(Supp. Figure S1D)**, consistent with a differentiated immunosuppressive macrophage phenotype engaged in extracellular matrix remodeling and clearance of dying cells. In post-CED IN samples, specifically, Mo-TAMs also showed enriched expression of *PDE3B* and *PTPN13*, suggesting inflammasome activation and M2 polarization. In contrast, Mg-TAM signatures in pre-CED biopsies were dominated by homeostatic microglial markers such as *TMEM119*, *P2RY12*, and *AQP4* **(Supp. Figure S1E)**, consistent with loss of a resting-state program after treatment.

We next stained matched pre- and post-CED infusion-zone biopsy slides for Iba1 (pan-myeloid marker), MSR1, MARCO, and pH2AX **(Figure 1G)**. Both MSR1 and MARCO indicate mesenchymal Mo-TAM states and have been previously associated with Mo-TAM enrichment, resistance to standard-of-care treatment, and worse survival in recurrent GBM (3, 5, 30–32). pH2AX is an early marker of double-stranded DNA breaks and serves as a proxy for topotecan-induced genotoxicity (33, 34). There was no significant reduction in overall cellularity within the drug infusion zone (Figure 1H) or reduction in labeling indices for Iba1+ and MSR1+ cells **(Figure 1I)**. By contrast, MARCO and pH2AX labeling indices were both significantly increased in the post-treatment infusion zone, showing 3.1- and 4.5-fold enrichment, respectively (patient-adjusted *p* = 0.002 and 0.038 for MARCO and pH2AX, respectively). Within Iba1+ myeloid cells, MSR1 positivity did not change significantly with treatment (**Figure 1J**). Rather, MSR1+ Iba1+ labeling indices showed substantial variability before treatment and more consistent high expression after treatment, suggesting consolidation of the myeloid phenotype in the drug infusion zone. MARCO and pH2AX were also significantly increased within the myeloid compartment (**Figure 1J**; 2.6- and 3.3-fold enrichment, patient-adjusted *p* = 0.001 and 0.018, respectively). Stratification by MSR1 status showed that MARCO induction was broadly distributed across both MSR1+ and MSR1− myeloid cells (**Supp. Figure S2A-B;** p = 0.035 and 0.004, respectively), whereas pH2AX increased significantly only in MSR1− myeloid cells (p = 0.004), with no significant change in MSR1+ cells. These findings suggest that MARCO-associated macrophage activation is distributed across MSR1-defined myeloid compartments, while persistent DNA-damage labeling after 28 days of infusion is most evident in MSR1− myeloid cells, potentially reflecting differential persistence, repair, or turnover across myeloid subsets.

### 3. CED-topotecan prolongs survival and induces myeloid DNA damage in a syngeneic PDGFα-driven murine glioma model

Because the human specimens capture a late treatment time point, we next used a syngeneic murine glioma model to examine earlier myeloid responses to topotecan. We performed catheter-delivered local topotecan treatment in a PDGFα-driven p53−/− murine glioma model using the same topotecan concentration (146 μM) as that used in the clinical trial (17). Dose-response studies confirmed that PDGFα-driven p53−/− murine glioma cells were sensitive to topotecan treatment *in vitro* (**Figure S3A**). As depicted in **Figure 2A**, tumor-bearing mice were randomized to receive either CED-topotecan or vehicle control beginning 21 days after glioma cell implantation. In separate survival and tissue-analysis cohorts, mice were either followed to humane clinical endpoints or sacrificed 3 or 7 days after initiation of pump-mediated infusion for downstream analysis. Successful intraparenchymal drug distribution was confirmed in all animals by MRI detection of co-infused gadolinium contrast (**Figure 2B**). CED-topotecan was well tolerated, with no signs of neurologic distress or treatment-related morbidity. Kaplan–Meier analysis showed a modest but significant survival benefit in topotecan-treated mice relative to controls (**Figure 2C**; median survival, 65 vs 45.5 days post-implantation; log-rank Mantel–Cox *p* < 0.0001), consistent with anti-proliferative activity within tumor cells. However, CED-topotecan was not curative in this model, as all mice reached humane endpoints before 90 days regardless of treatment, suggesting persistent or recurrent tumor activity and/or deleterious post-treatment inflammatory remodeling.

**Figure 2:**
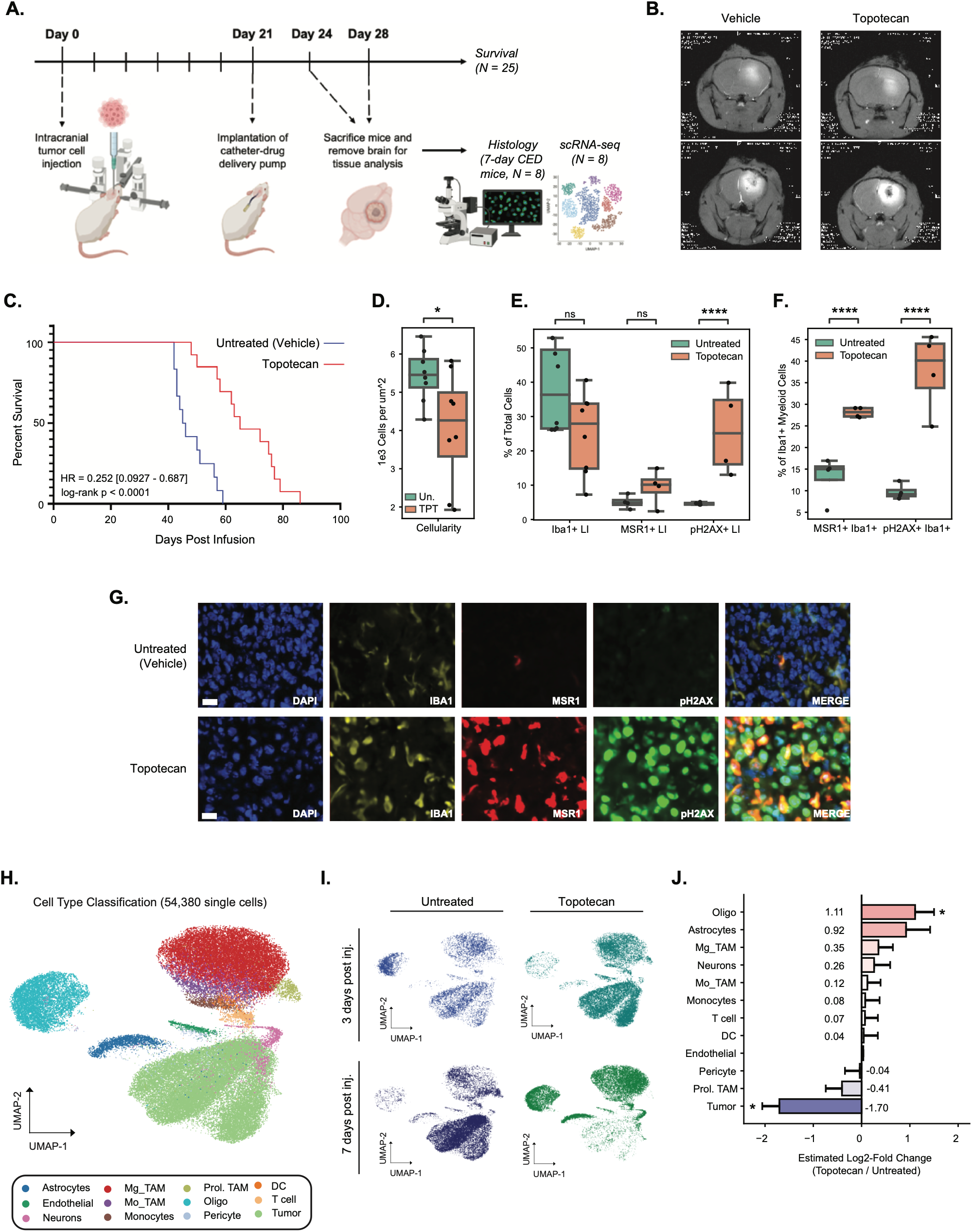
CED-topotecan prolongs survival but induces DNA damage and inflammatory myeloid reprogramming *in vivo*. **(A)** Experimental schematic of the PDGFα-driven syngeneic murine glioma model with CED of topotecan or vehicle control. Mice were stratified to survival analysis or tissue-based analysis at 3 or 7 days after initiation of pump-mediated infusion. **(B)** Representative MRI images from mice demonstrating the volume of gadolinium distribution in topotecan and vehicle conditions. **(C)** Kaplan–Meier survival analysis demonstrating significantly prolonged survival following CED-topotecan treatment. Median survival was 45.5 days for vehicle-control mice and 65 days for CED-topotecan-treated mice. Statistical significance was assessed using the log-rank Mantel–Cox test, and the hazard ratio was calculated using the log-rank method. **(D-F)** Quantification of cellularity (D), marker labeling indices (E), and within-myeloid positivity of MSR1 and pH2AX (F) following treatment. **(G)** Representative immunofluorescence staining for IBA1, MSR1 and pH2AX. Scale bar = 20 µm. **(H-I)** Single-cell RNA-sequencing UMAP visualization of major cellular populations in vehicle- and topotecan-treated tumors. **(J)** Differential abundance of overall cell types showing reduction of tumor cells and relative stability of the myeloid compartment after treatment. Tissue-based quantifications were analyzed using mixed linear-effects models with mouse-aware repeated-measures correction, accounting for two stained tissue sections analyzed per mouse brain, with n = 8 sections from n = 4 mice per condition. P values were adjusted for multiple comparisons using the Benjamini–Hochberg false discovery rate procedure. *BH-adjusted P < 0.05; ****BH-adjusted P < 0.0001. Differential abundance statistical markers represent FDR threshold for credible effects in scCoda < 0.05.

We next performed histological analysis of control and topotecan-treated mice after seven days of CED, staining for Iba1, MSR1, and pH2AX (**Figure 2G**). Overall cellularity was reduced 1.33-fold in topotecan-treated mice (**Figure 2D**; *p* = 0.035), consistent with tumor cell death. There were no significant changes in the proportional abundance of Iba1+ myeloid cells or overall MSR1 labeling **(Figure 2E)**. In contrast, pH2AX labeling was significantly increased following CED-topotecan by 5.55-fold, consistent with treatment-induced DNA damage **(Figure 2E**; *p* < 0.0001). Within the Iba1+ myeloid compartment, MSR1+ and pH2AX+ fractions increased by 1.88-fold and 4.37-fold, respectively, indicating phenotypic activation and DNA-damage signaling among surviving myeloid cells **(Figure 2F**; *p* < 0.0001 for each comparison). These findings indicate that seven days of CED-topotecan reduces tumor cellularity while inducing DNA damage and phenotypic remodeling within the surviving myeloid compartment. This model recapitulated the myeloid DNA damage observed in the CED-topotecan human cohort and prompted further analysis of the temporal dynamics of myeloid reorganization between 3 and 7 days.

### 4. scRNA-seq of 3- and 7-day-treated mouse tumors defines temporally structured inflammatory and DNA-damage programs in Mg-TAMs and Mo-TAMs following topotecan exposure

We performed 10X single-cell RNA sequencing (scRNA-seq) on untreated and CED-topotecan mouse tumors at early (3-day) and later (7-day) time points (n = 2 biological replicates per treatment condition and time point). After stringent filtering for mitochondrial reads, contaminating ribosomal transcripts, and predicted doublets, we recovered 54,380 single-cell events from 8 mice and identified tumor, myeloid, oligodendrocyte, neuron, endothelial, and T cell compartments. **Figure 2H-I** shows UMAP embeddings of cells from untreated and CED-topotecan mice, both combined and split by treatment and time point. Within the myeloid compartment, we resolved Mg-TAM, Mo-TAM, monocyte, and dendritic cell populations, with marker expression supporting these annotations **(Supp. Figure S3B)**. Mg-TAMs were enriched for *Hexb*, *P2ry12*, and *Tmem119*; Mo-TAMs for *Lyz2* and *Spp1*; and monocytes for *Ly6c2* and *Ccr2*.

We next assessed differential cell-type abundance across treatment and time point conditions using scCODA (**Figure 2J** and **Supp. Figure S3C**). Topotecan treatment was associated with a significant 3.3-fold depletion of tumor cells at 7 days (FDR *p* = 0.001), accompanied by a reciprocal 2.16-fold increase in oligodendrocyte abundance (FDR *p* = 0.025), likely reflecting replacement of tumor by white matter in these fixed-size biopsy specimens. No other major cell-type shifts reached statistical significance. Within the myeloid compartment, Mo-TAM, monocyte, and Mg-TAM proportions did not differ significantly between treated and untreated tumors at 7 days. Thus, compositional reorganization at 7 days was dominated by topotecan-induced tumor cell depletion, without major changes in myeloid abundance.

To examine myeloid-state dynamics in greater detail, we analyzed Mg-TAM and Mo-TAM populations separately. **Figure 3A** shows UMAP embeddings of each population, and **Figure 3B** shows expression of homeostatic markers (*P2ry12*, *Cx3cr1*, *Siglech*), activation markers (*Il1b*, *Lyz2*, *Clec7a*, *Cdkn1a*), and DNA damage/proliferation markers (*Trp53*, *Mki67*, *Top1/2a*), revealing heterogeneity within both populations. To characterize this heterogeneity in an unsupervised manner, we performed multi-omic factor analysis (MOFA) separately in Mg-TAMs and Mo-TAMs. In both populations, the top factor captured a polarized inflammatory module: in Mg-TAMs this was defined by *Cdkn1a*, *Clec7a*, *Lyz2*, *Ifrd1*, *Csf1r*, and *Il1a*, whereas in Mo-TAMs it was driven by *Prdx1*, *Lgals3*, *Cst7*, *Sepp1*, and *Hmox1* (**Supp. Figure S3D**). These findings indicate substantial intra-population heterogeneity across treatment conditions and time points.

**Figure 3:**
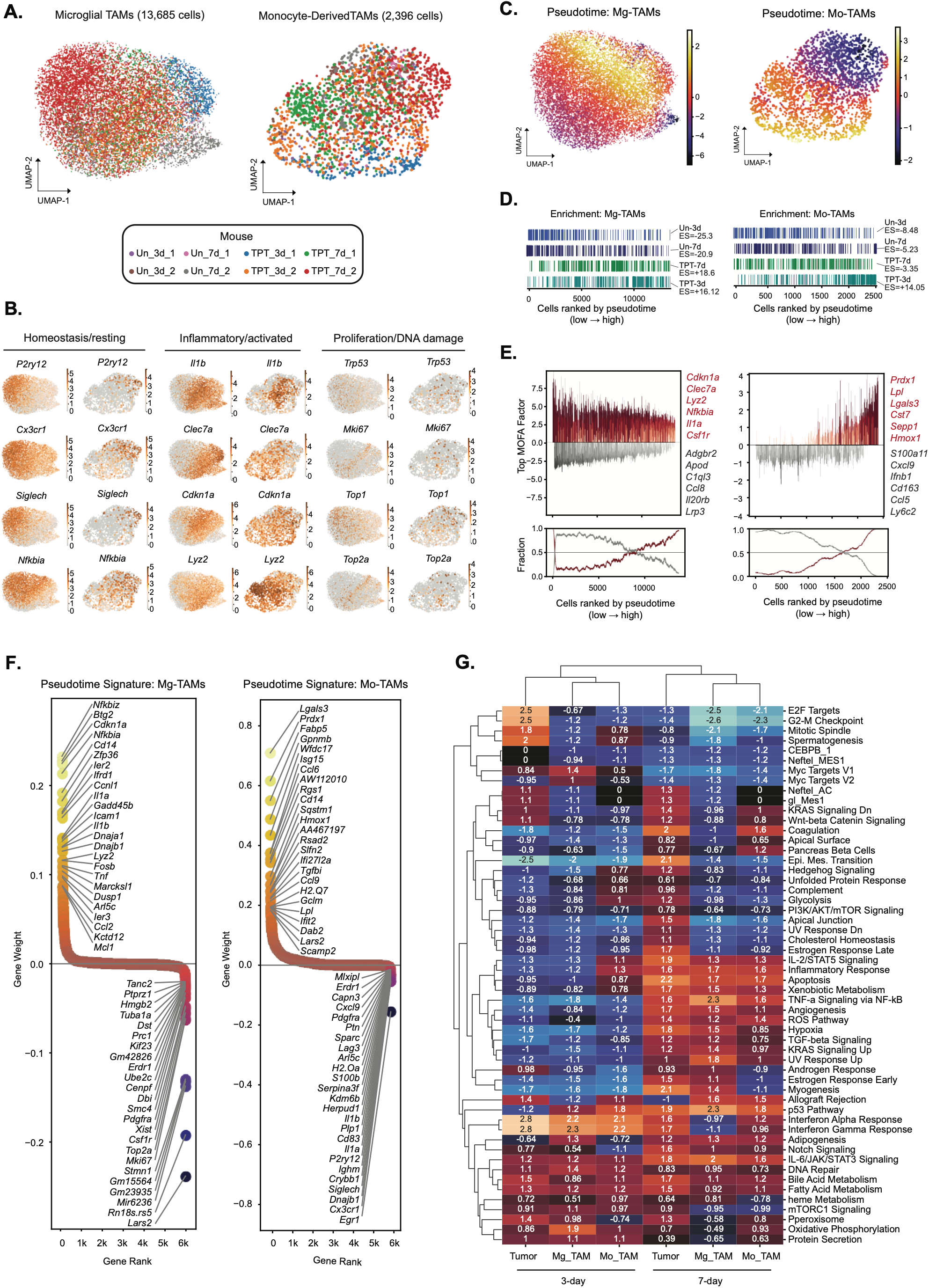
Single-cell RNA sequencing of murine gliomas captures myeloid transcriptional evolution following 3- and 7-day Topotecan exposure *in vivo*. **(A)** Single myeloid population UMAPs of the myeloid compartment, classified using label transfer from Pombo et al *Ccr2-*KO mice (1). **(B)** Single myeloid population UMAPs depicting heterogeneity of single-cell expression of homeostatic, inflammatory, and damage/proliferation markers. **(C)** UMAP embeddings showing cell-level pseudotime z-scores on single cell-type embeddings. **(D)** Rank-based statistical analysis of Mg-TAM, Mo-TAM, and tumor pseudotime trajectories, demonstrating that each recapitulates progression along ground-truth treatment/time condition annotations. Rank-based pseudotime enrichment across conditions was significant for each trajectory shown, with empirical p < 0.05. **(E)** Barcode enrichment plots (top) illustrating cell-level representation of MOFA scores across cells ranked by pseudotime (x-axis), with running fractional distributions of total signature-positive and signature-negative cells along pseudotime trajectory shown on the bottom of each plot. **(F)** Multi-omic factorization analysis (MOFA) of individual Mg-TAM and Mo-TAM populations was correlated with pseudotime scores to derive unsupervised pseudotime associated inflammatory signatures. **(G)** Gene set enrichment analysis of Hallmark pathways demonstrating activation of IL-1, TNF, interferon, and inflammatory response pathways in Mo-TAMs and Mg-TAMs from treated tumors, along with reduced markers of proliferation and increased DNA damage pathways.

We next inferred single-cell trajectories across treatments and time points and oriented pseudotime using untreated 3-day myeloid cells as the root state (Supplemental Methods). Resting microglia from peripheral non-neoplastic brain initially dominated Mg-TAM pseudotime analysis (**Supp. Figure S4**). To minimize this spatial sampling bias specifically for this analysis, we regressed a homeostasis signature (*P2ry12*, *Siglech*, *Tmem119*, *Cx3cr1*, etc.) from the Mg-TAM expression matrix prior to pseudotime inference, thus focusing on variation orthogonal to margin-versus-core differences. Cell-level pseudotime z-scores for each population are projected onto UMAPs embeddings in **Figure 3C**. To test the relationship between pseudotime and experimental condition, we used rank-based modeling to assess enrichment of each treatment/time point group across the ordered pseudotime axis, from low (t = 0) to high (t = 1). As shown in **Figure 3D**, pseudotime trajectories in Mg-TAM and Mo-TAM populations were both significantly associated with treatment and time (all rank-based enrichment *p* < 0.001). In both Mg-TAMs and Mo-TAMs, high pseudotime cells were enriched for topotecan-3d and topotecan-7d conditions, whereas untreated cells were concentrated at lower pseudotime values. High pseudotime Mo-TAMs were more enriched for topotecan-3d cells than topotecan-7d cells, suggesting a transient Mo-TAM inflammatory state early during treatment. To characterize these responses, we plotted enrichment of the top Mg-TAM and Mo-TAM MOFA factors across pseudotime (**Figure 3E**), which demonstrated strong enrichment of inflammatory modules at high pseudotime in both myeloid populations. These pseudotime-associated states were also linked to increased DNA damage response, proliferation, and mesenchymal and Mo-TAM identity scores (**Supp. Figure S3E**) (all rank-based enrichment *p* < 0.001). Finally, unsupervised gene signatures derived from MOFA scores and associated with increasing pseudotime (**Figure 3F**) identified strong population-specific inflammatory modules in Mg-TAMs and Mo-TAMs. In Mg-TAMs, high-pseudotime cells were enriched for NF-κB–associated and pro-inflammatory stress-response genes, including *Nfkbiz*, *Nfkbia*, *Cdkn1a*, *Il1a*, *Il1b*, *Tnf*, and *Ccl2*, whereas Mo-TAMs were marked by a distinct damage-associated macrophage program enriched for *Lgals3*, *Gpnmb*, *Fabp5*, *Hmox1*, *Isg15*, *Rsad2*, and *Tgfbi*. Together, these findings indicate that topotecan induces temporally structured inflammatory programs in both Mg-TAMs and Mo-TAMs.

Finally, we performed pseudo-bulk differential expression analysis comparing untreated and topotecan-treated cells within each population (Mg-TAM, Mo-TAM, tumor) at 3- and 7-day time points (**Figure 3G)**. In tumor cells, the 7-day response was marked by stronger downregulation of proliferative pathways versus control relative to the effect at 3 days, with reciprocal induction of cytokine production, DNA repair pathways (UV Response and p53), and stress- and microenvironment-remodeling programs (Epithelial/Mesenchymal Transition, Hypoxia/Apoptosis, Angiogenesis). Mg-TAMs and Mo-TAMs similarly exhibited decreased proliferative activity together with enrichment of inflammatory pathways, including IL-6 and TNFa signaling. By 7 days, both myeloid populations also showed greater enrichment of stress-response, hypoxia, angiogenesis, and TGF-β signaling programs. In contrast to pseudotime, which primarily captured a shared inflammatory trajectory across both 3-day and 7-day treated myeloid cells, pseudo-bulk analysis more clearly distinguished the later 7-day state, which showed stronger tissue-remodeling and mesenchymal-like features.

### 5. Acute ex vivo slice culture reveals microenvironmental influences on topotecan-induced myeloid responses

Having defined temporally structured myeloid remodeling in the murine CED model, we next asked how acute topotecan exposure affects human tumor and immune cells within a preserved tumor microenvironment. To examine how local cellular context shapes early human treatment responses, we treated patient-derived GBM slices with topotecan in an established acute *ex vivo* slice culture model **(Figure 4A)**. We performed microwell-based scRNA-seq on paired DMSO- and topotecan-treated slices from three newly profiled GBM patients and integrated these data with two previously published paired slice-culture datasets generated using the same experimental framework (35), yielding a combined cohort of five patient-derived GBM slice cultures. After stringent QC filtering, we recovered 71,616 single-cell events across 10 samples. **Figure 4B** shows a UMAP embedding of captured cells, color-coded by cell type (left) and split by treatment condition (right). **Figure 4C** shows the compositional distribution of each pair of slices, with predominance of tumor and myeloid populations. scCODA differential abundance analysis showed remodeling of the myeloid compartment after topotecan exposure, as Mo-TAMs decreased in proportional abundance by 2.8-fold (FDR-adjusted *p* = 0.003) (**Figure 4D**). Tumor cell proportional abundance remained stable after topotecan treatment (**Figure 4D**). However, MES-like tumor scores decreased nearly 2-fold across patient slices (**Figure 4F-G**; *p* = 0.013), whereas other tumor subtype scores did not change significantly. These findings indicate acute neoplastic-state rewiring without major loss of tumor representation.

**Figure 4:**
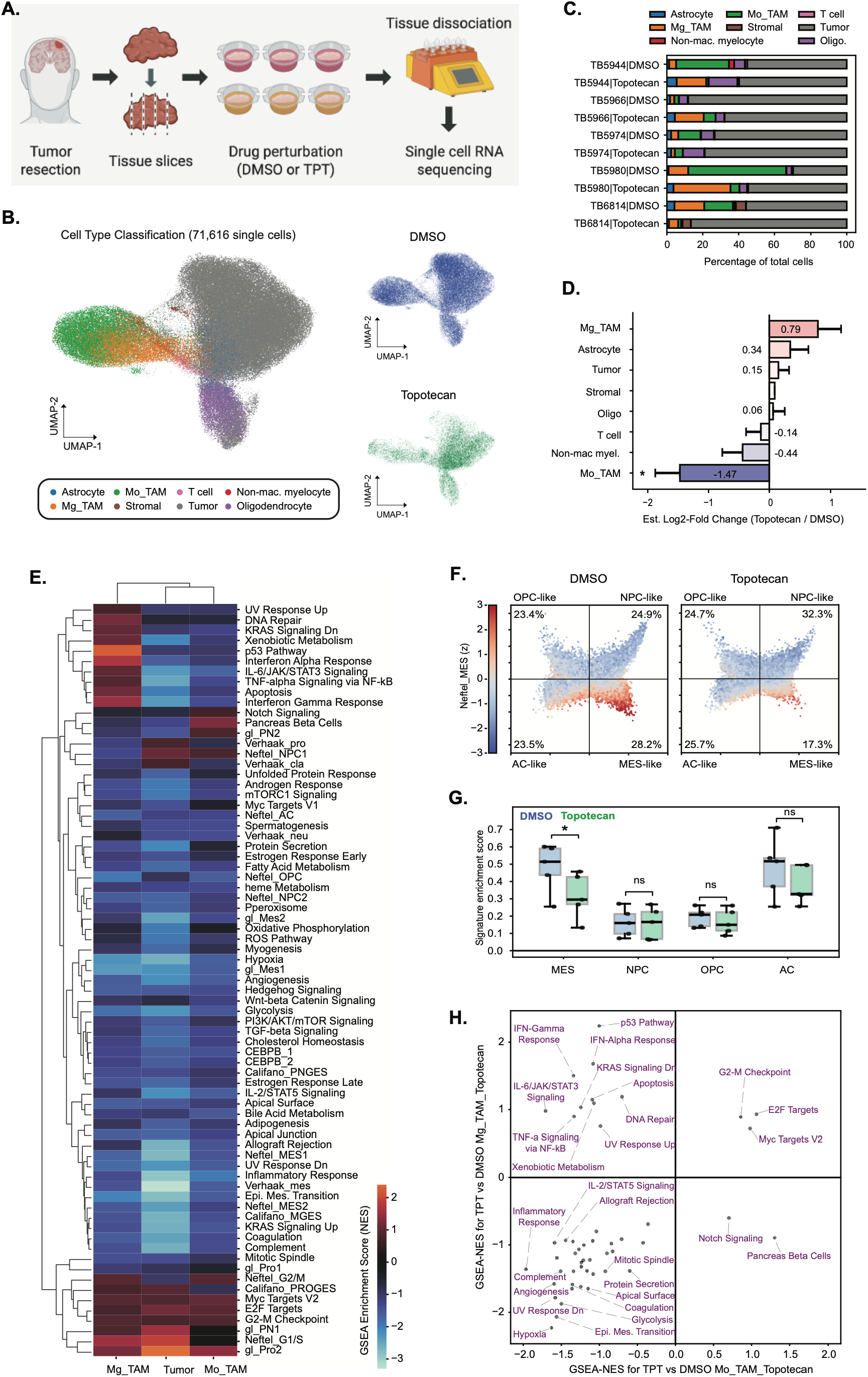
Acute treatment of ex vivo slice cultures with topotecan reveals microenvironmental influences on early microglial inflammatory responses. **(A)** Experimental schematic of the human GBM slice culture workflow, from tumor resection and generation of acute tissue slices through drug perturbation with topotecan or DMSO, tissue dissociation, and single-cell RNA sequencing. **(B)** UMAP visualization of integrated single-cell transcriptomes from human GBM slice cultures treated with topotecan or DMSO. (**C)** Total cell type proportional abundance profiles for each of 10 matched slices from *n =* 5 patients. **(D)** Differential abundance analysis demonstrating depletion of monocyte-derived macrophage states but preservation of microglia following topotecan exposure, with relatively stable tumor-cell abundance. Statistical markers represent FDR threshold for credible effects in scCoda < 0.05. **(E)** Heatmap depicting differential GSEA of Hallmark pathways within Mo-TAMs, Mg-TAMs, and Tumor cells post-acute topotecan, identifying differential proliferation modulation, inflammatory response and DNA-damage signaling across treatments and cell types. **(F-G)** Projection of single tumor-cell scores for Neftel OPC-like, NPC-like, AC-like, and MES-like signatures and score quantitation by slice. **(H)** Quadrant plot comparing Hallmark GSEA normalized enrichment scores (NES) from topotecan versus DMSO differential expression signatures in Mg-TAMs and Mo-TAMs, highlighting concordant and discordant pathway responses between myeloid populations in slice culture.

Next, we performed sample-wise pseudo-bulk differential expression analysis within tumor, Mo-TAM, and Mg-TAM populations, followed by Hallmark GSEA (**Figure 4E**). In tumor cells and Mo-TAMs, enrichment scores were broadly suppressed, with only minor enrichment of cell cycle/proliferation pathways (E2F Targets, Myc Targets V2, G2-M Checkpoint), consistent with widespread genotoxic stress and impending cell death. By contrast, Mg-TAMs showed marked enrichment of DNA damage and stress response pathways (DNA Repair, UV Response, p53 pathway, Apoptosis), as well as increased proliferation, similar to Mo-TAMs and tumor cells. Mg-TAMs were also enriched for inflammatory pathways, including IFN-Alpha/Gamma, IL-6/JAK/STAT3, and TNFa, whereas mesenchymal/ECM remodeling pathways were strongly downregulated following topotecan. Direct comparison of Hallmark GSEA normalized enrichment scores from topotecan versus DMSO signatures in Mg-TAMs and Mo-TAMs further showed that inflammatory and DNA damage-stress pathways were more strongly enriched in Mg-TAMs, whereas Mo-TAMs retained some proliferative capacity and showed little direct inflammatory induction by topotecan **(Figure 4H)**. These findings suggest that the acute slice-culture model captures an early Mg-TAM–skewed inflammatory and DNA-damage response that may precede the broader myeloid remodeling observed in post-CED clinical biopsies.

### 6. Topotecan induces a stress-coupled inflammatory program in human microglia and enhances macrophage phagocytosis in vitro

While slice culture preserves tumor–immune architecture, it does not fully disentangle direct drug effects from microenvironmentally mediated responses. We therefore used isolated and co-culture systems to define direct topotecan-induced microglial inflammatory programs and tumor–macrophage interactions. First, we cultured induced pluripotent stem cell-derived microglia (iPSC-Mg) with 2.5 μM topotecan for 2 days and measured transcriptomic changes relative to DMSO-treated controls by bulk RNA-seq. In parallel, we treated iPSC-Mg with lipopolysaccharide (LPS) as a positive control to compare topotecan-induced inflammatory responses with a more canonical activation stimulus. We used iPSC-Mg rather than immortalized cell lines because they more closely resemble homeostatic human microglia.

**Figures 5A-B** show volcano plots of the most significantly differentially expressed genes in LPS- and topotecan-treated iPSC-derived microglia relative to control, respectively. Topotecan upregulated stress-response and immune activation genes, including *NUPR1*, *G0S2*, and *TREM1*, while downregulating canonical cell cycle regulators, including *MYBL2*, *CDK1*, *KIF2C*, and *KIF14*, consistent with suppression of proliferative and mitotic programs. By contrast, LPS preferentially induced classical chemokine and interferon-responsive genes, including *CXCL8*, *CCL3*, *IFIT3*, and *OAS2*, without the same coordinated suppression of proliferative pathways. Gene set enrichment analysis using MSigDB Hallmark terms demonstrated preferential induction of damage-associated inflammatory pathways by topotecan relative to LPS, including IL-6/JAK/STAT3, IL-2/STAT5, UV Response, Hypoxia, and Complement, together with stronger suppression of Apoptosis and proliferative pathways, including Myc Targets V1/V2 and E2F Targets (**Figure 5C**). LPS induced stronger canonical inflammatory pathways, including TNF-alpha and IFN-Alpha/Gamma signaling. These results demonstrate that topotecan induces a distinct stress-coupled inflammatory state in microglia compared with canonical LPS activation.

**Figure 5:**
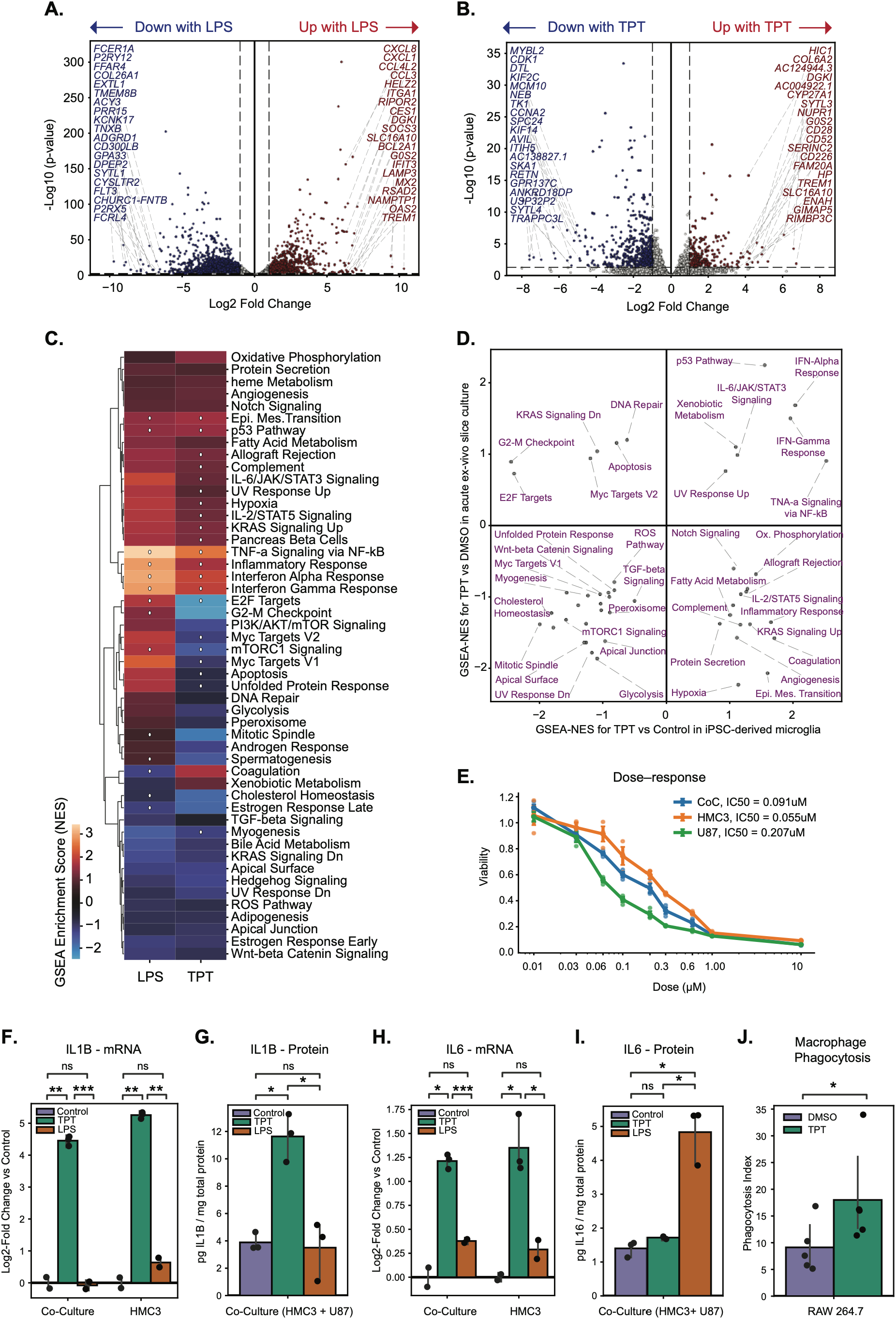
Acute topotecan treatment *in vitro* defining direct inflammatory and DNA-damage responses in cultured microglia. **(A-B)** Volcano plots showing differential expression signatures and top DE genes between iPSC-Mg treated *in vitro* with LPS (A) and topotecan (B) vs DMSO control. **(C)** Heatmap depicting Hallmark gene set enrichment scores in the LPS and topotecan signatures. (D) Quadrant plot comparing Hallmark GSEA normalized enrichment scores (NES) from topotecan versus DMSO differential expression signatures in Mg-TAMs in slice culture and iPSC-derived microglia, highlighting concordant and discordant pathway responses between myeloid populations in slice culture. **(E)** Dose-response curves to topotecan for both HMC3 microglia and U87 glioma cell lines alone and in co-culture, measured as normalized viability. **(F-I)** Inflammatory cytokine responses in monocultured and co-cultured HMC3 microglia following exposure to topotecan or LPS. IL1B (F) and IL6 (H) mRNA expression are shown as log2 fold change by qPCR, while IL1B (G) and IL6 (I) protein levels were quantified by MSD assay. **(J)** Functional assay measuring phagocytosis of topotecan-treated p53−/− PDGFA cells by murine bone-marrow derived myeloid cells (RAW 264.7), distinguished by fluorescence-activated cell sorting. Statistics were calculated using Student’s t-test with pairing as appropriate and ANOVA with Tukey’s post-hoc test for multiple comparisons. (*) represents *p* < 0.05; (**) represents *p* < 0.01.

Next, we compared the Mg-TAM slice culture signature with the corresponding response in isolated iPSC-derived microglia. We plotted Hallmark GSEA normalized enrichment scores from topotecan versus DMSO differential expression signatures in slice-culture Mg-TAMs against those from topotecan-treated iPSC-Mg (**Figure 5D)**. This comparison showed preservation of many damage-response and inflammatory pathways across systems (**Figure 5D**, top right quadrant), including IFN-Alpha/Gamma, IL-6/JAK/STAT3, TNFa, p53, and UV Response. Additional proliferative and DNA-damage pathways (DNA Repair, Apoptosis, G2-M Checkpoint, E2F and Myc Targets) were selectively enriched in Mg-TAMs within the preserved slice-culture microenvironment, whereas MES-inflammatory pathways, including Hypoxia, Epithelial-Mesenchymal Transition, Coagulation, Complement, and IL-2/STAT5 were selectively suppressed. Together, these data suggest that acute topotecan exposure induces a conserved microglial damage/inflammatory response that is further shaped by the preserved slice-culture microenvironment, without recapitulating the chronic Mo-TAM/MARCO-like state observed in chronically treated human biopsy tissue.

We next treated human HMC3 microglial cell lines with topotecan for 3 days, either alone or in co-culture with U87 glioma cells. Dose response studies demonstrated dose-dependent viability effects in both cell lines alone and in co-culture, supporting 100 nM topotecan as a suitable treatment dose because it exceeded the IC_50_ of U87 glioma cells while remaining below the IC_50_ of HMC3 microglia (**Figure 5E**). *IL1B* and *IL6* mRNA expression were measured by quantitative PCR of cell lysates and plotted as log2 fold change relative to control **(Figures 5F and 5H)**.

Topotecan induced *IL1B* and *IL6* mRNA expression in both HMC3 monoculture and HMC3-U87 co-culture (*p =* 0.0096 and 0.007, respectively, for *IL1B; p =* 0.014 and 0.025, respectively, for *IL6*), whereas LPS-induced changes in *IL1B* and *IL6* mRNA were not significant in either condition (monoculture and co-culture: p = 0.104 and 0.745 for *IL1B*; p = 0.185 and 0.155 for *IL6*). Similar induction across monoculture and co-culture conditions suggests that this response does not require co-culture-dependent cues. We also used MSD assays to quantify IL1B and IL6 protein levels in co-culture lysates (**Figures 5G** and **5I**, respectively). IL1B protein levels were increased following topotecan treatment compared with both control and LPS-treated samples (*p =* 0.0096 and 0.008, respectively), whereas LPS did not significantly increase IL1B protein levels compared with control (*p* = 0.791). In contrast, IL6 protein levels increased following LPS treatment (*p* = 0.015), but not topotecan treatment (*p* = 0.138). Together, these *in vitro* experiments indicate that topotecan is sufficient to induce a stress-coupled inflammatory transcriptional response in human microglia, with corresponding protein-level induction of IL1B.

Finally, to assess whether topotecan also modulates tumor-myeloid interactions through tumor-intrinsic changes, we performed a phagocytosis assay in which tumor cells were pre-treated with topotecan (5 µM, 72 hours), washed, and then co-cultured with RAW 264.7 macrophages. Macrophages exhibited 1.97-fold increased phagocytosis index of topotecan-treated tumor cells compared with untreated controls (*p* = 0.014) (**Figure 5J**). These results suggest that topotecan-induced changes in tumor cells increase susceptibility to macrophage phagocytosis, potentially shaping tumor– myeloid interactions within the treatment-affected microenvironment.

## DISCUSSION

In this study, we characterized the spatial, cellular, and temporal features of the early response to local chemotherapy in GBM. In addition to suppressing proliferative neoplastic cells, topotecan induced genotoxic stress and inflammatory transcriptional programs within tumor-associated myeloid populations. By integrating paired MRI-localized human biopsies with murine tumors analyzed at 3- and 7-day treatment time points, human slice cultures, and isolated human microglial models, we demonstrate that the response to local chemotherapy extends beyond tumor cell injury to encompass inflammatory remodeling of myeloid cells. Together, these findings define the early cellular and molecular events associated with therapy-associated inflammatory remodeling of the GBM microenvironment.

A central feature of this study was comparing pretreatment biopsies with immediate post-treatment biopsies from the same patients, while also localizing post-treatment specimens relative to the maximal infusion zone. This design allowed us to examine proximal chemotherapy-associated tissue changes before clinical or radiographic recurrence, rather than relying on recurrent specimens that reflect the cumulative effects of therapy, tumor regrowth, and longitudinal evolution (18, 21, 22). Because each patient served as their own pretreatment control and post-treatment specimens were stratified by proximity to the MRI-defined infusion zone, the clinical biopsy analysis provided spatially localized evidence of treatment-associated transcriptional remodeling. By combining this clinical sampling strategy with 3-day and 7-day murine CED models, acute human slice culture, and isolated human microglial systems, our approach enabled characterization across complementary temporal and cellular contexts.

In human tumors, transcriptional remodeling was most pronounced within the infusion zone, with suppression of proliferative tumor programs and enrichment of inflammatory cytokine signaling, interferon responses, hypoxia adaptation, and mesenchymal features. This spatial pattern suggests that the inflammatory and mesenchymal transcriptional state reflects a localized response associated with pharmacologic exposure, superimposed on broader tumor evolution. Cell-type-resolved analyses further linked this shift to myeloid remodeling, including enrichment of monocyte-derived macrophages and induction of a MARCO-positive phenotype. Multiplex immunofluorescence confirmed expansion of MARCO-positive myeloid populations, and pH2AX staining demonstrated treatment-associated DNA damage in Iba1-positive myeloid cells within the infusion zone. Together, these findings indicate that immediately after 28 days of treatment, the infusion zone is characterized by spatially localized myeloid remodeling, including macrophage enrichment, MARCO-positive myeloid populations, and genotoxic stress within tumor-associated myeloid cells.

Murine modeling clarified earlier temporal features of CED-topotecan-induced myeloid remodeling. Single-cell RNA sequencing demonstrated broad inflammatory activation within the myeloid compartment, with induction of interferon, IL-6/JAK/STAT3, and TNF-α/NF-κB signaling in both Mg-TAMs and Mo-TAMs, alongside DNA-damage responses and suppression of proliferative programs. Pseudotime analysis placed both 3-day and 7-day treated myeloid cells along a shared inflammatory trajectory relative to untreated cells, indicating that inflammatory activation is a common feature across treated states. By contrast, pseudo-bulk comparison more clearly resolved the later 7-day state, which showed stronger hypoxia, angiogenic, TGF-β, tissue-remodeling, and mesenchymal-like programs. Notably, these transcriptional changes occurred without increased overall myeloid abundance by histology. Instead, increased DNA damage in tumor-associated myeloid cells and increased MSR1 expression suggest that the post-treatment myeloid compartment undergoes phenotypic remodeling in association with local genotoxic stress, although contributions from altered recruitment, survival, or turnover cannot be excluded.

Human slice culture and *in vitro* systems further clarified the relationship between preserved microenvironmental context and isolated microglial responses to topotecan exposure. Acute patient-derived slice cultures preserved tumor– immune architecture and showed that Mg-TAMs mount prominent DNA-damage and inflammatory responses after topotecan exposure. In this model, topotecan exposure was also associated with decreased proportional abundance of Mo-TAMs and non-macrophage myelocytes, with reciprocal enrichment of Mg-TAMs. This pattern is consistent with differential sensitivity or persistence among myeloid subsets after acute topotecan exposure, although altered survival, proliferation, or state-dependent recovery from dissociation cannot be distinguished in this system. In parallel, iPSC-derived microglia and HMC3 cells demonstrated that topotecan can induce stress-coupled inflammatory signaling in human microglial models in isolation. Both the acute slice-culture response and the in vitro findings differed from the 28-day human biopsy response, which showed stronger monocyte-derived macrophage enrichment and MARCO-associated mesenchymal remodeling within the infusion zone. This contrast suggests that the acute response to topotecan is distinct from the 28-day treatment time point captured after sustained local exposure.

Together, these data support a model in which chemotherapy exposure produces parallel cytotoxic and microenvironmental effects in GBM, suppressing proliferative neoplastic programs while inducing genotoxic stress and inflammatory activation within tumor-associated myeloid cells. Across model systems, the response appears temporally structured: acute models highlight microglial inflammatory activation and DNA-damage-associated programs, whereas the tissue state captured immediately after 28 days of local exposure shows more prominent macrophage-associated mesenchymal remodeling and enrichment of epithelial–mesenchymal transition, hypoxia adaptation, and angiogenic pathways. Although these findings do not establish that therapy-induced myeloid remodeling directly drives recurrence or treatment resistance, they characterize cellular and molecular changes that arise during the post-treatment interval and may precede the later inflammatory microenvironment observed in recurrent disease (3, 5, 30, 36, 37).

Our findings generate testable hypotheses for future therapeutic design. First, they suggest that local chemotherapy should be evaluated not only by its cytotoxic effects on tumor cells, but also by its effects on the immune microenvironment. Second, the observation that myeloid inflammatory activation occurs at early treatment time points raises the possibility that timing may be important for future combination strategies. Interventions targeting acute inflammatory signaling, including IL-1, TNF, and interferon-associated pathways, may require different scheduling than approaches directed at later macrophage-associated tissue-remodeling programs. Emerging therapies targeting transcriptional regulators of mesenchymal and inflammatory macrophage programs may be particularly relevant (37, 38). Similarly, targeted anti-inflammatory therapies such as anakinra or broader immunosuppressants including dexamethasone could be explored as strategies to modulate treatment-associated myeloid remodeling after local chemotherapy (37, 39, 40). Patient-derived slice-cultures and murine CED models may provide useful platforms for evaluating how dose, duration, and treatment schedule shape the balance between tumor cytotoxicity and inflammatory remodeling (36). Additionally, our model of post-treatment myeloid dynamics provides a preliminary framework for interpreting inflammatory changes in prior and future intratumoral therapy studies.

Several limitations warrant consideration. The clinical cohort was constrained by the first-in-human design of the trial and the inherent scarcity of paired pre- and post-treatment tissue from an implantable pump-catheter procedure. The paired biopsy design provides biological resolution that does not depend on large patient numbers, but the generalizability of the specific myeloid remodeling programs identified here to broader recurrent GBM populations and other CED agents remains to be established. Temporal ordering was inferred from cross-sectional time points and trajectory analysis rather than direct lineage tracing. Differences between murine and human immune biology remain an inherent limitation, and lineage relationships among macrophage subsets cannot be resolved definitively using transcriptomics alone. Finally, while convergence across spatial human analyses, *in vivo* modeling, slice culture, and isolated microglial systems supports a treatment-associated myeloid response, the downstream consequences of these inflammatory states for recurrence and therapeutic resistance will require direct functional testing.

In summary, this study demonstrates that local chemotherapy exposure initiates a spatially structured and temporally evolving inflammatory myeloid response in GBM, linked to genotoxic stress in both neoplastic and immune compartments. By profiling paired, spatially annotated human biopsies alongside complementary experimental models, we define the cellular and molecular architecture of the post-chemotherapy microenvironment across multiple treatment time points. This multiparametric framework may also help prioritize treatment-associated cellular programs and molecular pathways for future therapeutic investigation. These findings highlight therapy-associated myeloid remodeling as a structured component of the local chemotherapy response and provide a foundation for future studies evaluating how treatment dose, schedule, and rational combinations shape the post-treatment tumor-immune microenvironment.

## METHODS

### Sex as a biological variable

Samples from the CED-topotecan clinical trial (17) were from both male and female patients. Orthotopic mouse experiments were conducted exclusively in female mice; the applicability of these findings to male mice remains unknown.

### Patient samples and clinical biopsy annotation

Recurrent GBM specimens were obtained from patients enrolled in a previously reported investigator-initiated phase 1b trial of CED-topotecan (NCT03154996) (17). Patients underwent stereotactic catheter implantation, 28 days of locally infused topotecan with co-infused gadolinium for MRI-based visualization of drug distribution, and catheter explant with post-treatment tissue sampling. Paired pre-CED and post-CED biopsies from five patients were analyzed. Pre-CED biopsies were defined as those obtained at catheter implantation, and post-CED biopsies were defined as those obtained at catheter explantation and were annotated as within or outside the maximal infusion zone based on intraoperative biopsy coordinates and post-infusion gadolinium-enhanced MRI localization, as previously described.

### Human biopsy immunofluorescence

Formalin-fixed paraffin-embedded biopsy sections from pre-CED and post-CED infusion-zone tissue were stained for myeloid and DNA-damage markers using antibodies against Iba1, MSR1, MARCO, and pH2AX, with DAPI nuclear counterstaining. Sections were imaged by confocal microscopy, and marker-positive cells were quantified from digitized images using standardized image-analysis workflows. Total cellularity, total marker labeling indices, and marker positivity within Iba1-positive myeloid cells were compared between pre-treatment and post-treatment infusion-zone specimens. Antibody sources, dilutions, antigen retrieval conditions, imaging parameters, and quantification details are provided in the Supplementary Methods.

### Bulk RNA-seq processing, differential expression, and pathway analysis

Raw aligned bulk RNA-seq counts from paired clinical biopsies were obtained from the original CED-topotecan trial dataset and aligned with biopsy-level clinical and spatial metadata. Genes corresponding to mitochondrial, ribosomal, hemoglobin, and non-coding features were removed before downstream analysis. For visualization and signature scoring, counts were library-size normalized and log transformed. Differential expression was performed on raw count matrices using PyDESeq2 after filtering low-count genes. Pairwise contrasts were generated among pre-CED, post-CED IN, and post-CED OUT biopsies, and gene-wise p values were adjusted using the Benjamini-Hochberg procedure. Differential expression signatures were visualized using volcano plots and infusion-zone quadrant plots comparing post-CED IN versus pre-CED and post-CED OUT versus pre-CED log2 fold-change values.

Gene-set enrichment analysis was performed on ranked differential expression signatures using the MSigDB Hallmark collection. Canonical GBM subtype signatures from Verhaak et al. (REF) were scored in each biopsy using normalized log-transformed expression, and subtype scores were compared across pre-CED, post-CED IN, and post-CED OUT groups (41). Detailed gene filtering, ranking, enrichment, multiple-testing correction, and visualization procedures are provided in the Supplementary Methods.

### Bulk deconvolution and high-resolution cell-type expression inference

Cell-type fractions in clinical bulk RNA-seq samples were inferred using CIBERSORTx with GBmap, a recently published single-cell RNA-seq atlas of glioblastoma comprising more than 115 patients, as the reference dataset (29, 42). To better match the treatment-exposed recurrent GBM cohort, the reference was restricted to recurrent biopsies only, yielding 35 patients across five studies while preserving sufficient cell numbers for deconvolution. The recurrent GBM reference was derived from annotated GBmap cell labels and collapsed into cell-type expression profiles for the major tumor and non-neoplastic compartments used in the study. Inferred cell-type fractions were compared across pre-CED, post-CED IN, and post-CED OUT samples.

CIBERSORTx high-resolution mode was additionally used to infer sample-wise cell-type-specific expression profiles, focusing on tumor, Mo-TAM, and Mg-TAM compartments. Cell-type-specific expression profiles were analyzed by differential expression and Hallmark enrichment as described above. Full reference construction, CIBERSORTx settings, and high-resolution analysis details are provided in the Supplementary Methods.

### Murine CED-Topotecan model, survival analysis, and tissue processing

Orthotopic tumors were generated in female C57BL/6 mice using a syngeneic PDGFA-driven, p53-deficient glioma model (43). At 21 days after tumor implantation, mice received catheter-based intraparenchymal delivery of matched vehicle or topotecan using subcutaneously implanted osmotic pumps (44, 45). Drug distribution was confirmed by MRI visualization of co-infused gadolinium. For survival experiments, mice were monitored until death or predefined humane endpoint, and survival was analyzed using Kaplan-Meier curves and the log-rank test. For tissue analyses, brains were collected at defined time points after initiation of pump-mediated infusion, fixed, embedded, sectioned, and processed for immunofluorescence or dissociated for single-cell RNA-seq. Additional surgical, dosing, imaging, tissue-processing, and animal-number details are provided in the Supplementary Methods.

### Murine immunofluorescence and image quantification

Mouse brain sections were stained for Iba1, MSR1, and pH2AX with DAPI counterstaining. Images were acquired by confocal microscopy and quantified using QuPath within standardized regions of interest centered on the catheter tract (46). Regions of interest were drawn blinded to treatment group. Total cellularity, Iba1-positive myeloid abundance, MSR1 labeling, pH2AX labeling, and marker positivity within Iba1-positive cells were quantified and compared between vehicle- and topotecan-treated tumors. Antibody details, image-analysis workflow, and statistical procedures are provided in the Supplementary Methods.

### Single-cell RNA-seq preprocessing, annotation, and differential abundance analysis

Mouse tumor and patient-derived slice-culture single-cell RNA-seq datasets were processed using standardized Scanpy-based workflows. Count matrices were filtered to remove low-quality cells, high-mitochondrial or high-ribosomal events, hemoglobin-rich events, low-detected-gene cells, predicted doublets, and confounding mitochondrial, ribosomal, hemoglobin, heat-shock, and non-coding genes. Counts were normalized, log transformed, and integrated using highly variable gene selection, principal component analysis, Harmony correction, k-nearest-neighbor graph construction, UMAP embedding, and Leiden clustering. Cell types were assigned using marker-based annotation and hierarchical label-transfer approaches from GBM single-cell references, with myeloid subsets further resolved using reference annotations distinguishing Mo-TAM and Mg-TAM states. For the murine CED-topotecan dataset, Mg-TAM and Mo-TAM annotations were assigned using label transfer from the Pombo Antunes et al. reference atlas (1), which includes CCR2-KO scRNA-seq data to support discrimination of resident microglia-derived and monocyte-derived tumor-associated myeloid populations. Differential abundance across treatment and time point groups was assessed using scCODA. Detailed preprocessing thresholds, reference datasets, annotation hierarchies, and model settings are provided in the Supplementary Methods.

### Murine myeloid factorization, pseudotime, pseudo-bulk differential expression, and pathway analysis

To characterize treatment-associated myeloid-state remodeling in the murine CED model, Mg-TAM and Mo-TAM populations were analyzed separately. Multi-Omics Factor Analysis (MOFA) was used to identify continuous transcriptional programs within each myeloid compartment (47). Diffusion-map and diffusion-pseudotime analyses were performed on integrated single-cell embeddings to order myeloid cells along treatment-associated transcriptional trajectories. Associations between pseudotime, treatment condition, time point, MOFA factors, and gene programs were quantified to derive myeloid pseudotime-associated signatures. In parallel, pseudo-bulk differential expression was performed within Tumor, Mg-TAM, and Mo-TAM populations by aggregating cells at the biological-sample level before differential expression and Hallmark enrichment analysis. Full MOFA, pseudotime, pseudo-bulk, and enrichment procedures are described in the Supplementary Methods.

### Patient-derived slice culture and single-cell perturbation analysis

Freshly resected human GBM tissue was processed using an established acute ex vivo slice-culture system (33). For this study, paired topotecan- and vehicle-treated slice-culture scRNA-seq data were generated from three newly processed GBM specimens and integrated with two previously published paired slice-culture datasets generated using the same experimental framework (35). Tissue slices were equilibrated on membrane inserts and treated with topotecan or vehicle control for 18 hours before single-cell RNA-seq processing. Additional culture conditions, dosing, and tissue-processing details are provided in the Supplementary Methods.

Single-cell profiles from paired treated and control slices were annotated into tumor and non-neoplastic compartments, with myeloid populations separated into Mo-TAM and Mg-TAM states. Treatment-associated compositional changes were analyzed using scCODA. Within-cell-type pseudo-bulk differential expression and Hallmark enrichment analyses were performed for Tumor, Mo-TAM, and Mg-TAM populations. Tumor-state scores and cross-system comparisons between slice-culture myeloid responses and *in vitro* microglial responses were calculated as described in the Supplementary Methods (48).

### In vitro microglial, glioma co-culture, cytokine, and phagocytosis assays

Human iPSC-derived microglia, HMC3 microglia, U87 glioma cells, and macrophage-based phagocytosis systems were used to evaluate direct and tumor-mediated effects of topotecan on inflammatory signaling and tumor-myeloid interactions (49–51). iPSC-derived microglia were treated with topotecan, lipopolysaccharide, or vehicle and analyzed by bulk RNA-seq and Hallmark enrichment. HMC3 microglia were treated in monoculture or U87 glioma co-culture, followed by quantitative PCR measurement of inflammatory cytokine transcripts and immunoassay-based quantification of cytokine protein levels. To assess treatment-induced changes in tumor-cell phagocytosis, murine glioma cells were pre-treated with topotecan or vehicle, washed, co-cultured with RAW 264.7 macrophages, and quantified for macrophage uptake. Cell-culture conditions, doses, time points, assay reagents, normalization procedures, and statistical tests are provided in the Supplementary Methods.

### Statistical analysis and reproducibility

Unless otherwise specified, statistical analyses were performed in Python using Scanpy, PyDESeq2, gseapy, scCODA, MOFA/muon, Harmony, SciPy, statsmodels, and related scientific-computing packages. Multiple-testing correction was performed using the Benjamini-Hochberg procedure where appropriate. Survival distributions were compared using the log-rank test. Boxplots and bar plots show the summary statistics indicated in the corresponding figure legends, and individual points represent the relevant experimental unit for each assay. Detailed software versions, analysis parameters, experimental units, and statistical frameworks are provided in the Supplementary Methods. A detailed sensitivity analysis using alternative analytical strategies—including patient-condition aggregation followed by ANOVA with Tukey’s post hoc tests, and Harmony adjustment for patient-associated variation followed by ANOVA with Tukey’s post hoc tests—is presented in **Supplementary Figure S1G–I** and **Supplementary Table 1**. These analyses were performed to assess whether the direction and magnitude of the observed human CED biopsy differences were robust to alternative approaches for handling patient-associated variation and the presence of multiple biopsies per patient.

### Study approval

All patients in the CED-topotecan trial provided written informed consent under a protocol approved by the Columbia University Institutional Review Board and the FDA. Additional clinical eligibility criteria and regulatory details are provided in the Supplementary Methods. All mouse experiments were approved by the Columbia University Institutional Animal Care and Use Committee and performed in accordance with institutional guidelines.

### Data availability

All new sequencing data and code will be made publicly available before publication on appropriate repositories (NCBI GEO for bulk RNA-seq and scRNA-seq; GitHub for code notebooks). Values for all data points in graphs are reported in the Supporting Data Values file. All relevant materials are available with reasonable request to the corresponding author.

## Supporting information

Supplemental Table 1 Statistics

Supporting Data Values

## AUTHORSHIP CONTRIBUTIONS

AJT, MRW, PJC, and DET are co-first authors. Authorship order was agreed upon after discussion among all co-first and co-senior authors. Experiments and analyses were designed by AJT, MRW, PJC, DET, NBD, AM, LL, PC, and JNB. Experiments were performed by AJT, PJC, DET, NBD, AM, and LL. Computational analyses were performed by MRW, with input from AJT, PJC, LL, PC, and JNB. SW provided statistical guidance and review. Figures were prepared by AJT, MRW, PJC, and DET. AJT, MRW, PJC, and DET wrote the initial draft of the manuscript, with input from LL, PC, and JNB. NBD, AM, MA, NWR, MGA, CPS, AXC, NI, AV, MK, AD, BP, AB, AK, NH, CS, JF, CK, WZ, ZL, HH, NK, ARA, NJW, NY, JAN, KRS, BJAG, PS, OAD, and JG contributed to experimental design, data collection, data analysis, interpretation of findings, and/or critical revision of the manuscript. LL, PC, and JNB provided study supervision, scientific guidance, interpretation of findings, and critical revision of the manuscript. PC and JNB acquired funding and provided overall oversight. All authors reviewed and approved the final manuscript.

## ACKNOWLEDGEMENTS

We thank the staff of the Molecular Pathology Shared Resource at the Herbert Irving Comprehensive Cancer Center at Columbia University Irving Medical Center for assistance with tissue embedding and sectioning. These studies also used the Confocal and Specialized Microscopy Shared Resource of the Herbert Irving Comprehensive Cancer Center at Columbia University, supported in part by the NIH/NCI Cancer Center Support Grant P30CA013696. Mouse MRI studies were performed in the MR Facility of the Oncology Precision Therapeutics and Imaging Core (OPTIC) Shared Resource, which is supported by the Columbia University Irving Medical Center Cancer Center Support Grant and NIH/NCI grant P30CA013696. Statistical support was provided by the Cancer Biostatistics Shared Resource of the Herbert Irving Comprehensive Cancer Center, supported in part by NIH/NCI Cancer Center Support Grant P30CA013696.

