## Supplemental Figures and Methods for "Paired Tumor Biopsies Reveal Spatiotemporal Myeloid Remodeling After Local Chemotherapy in Glioblastoma"

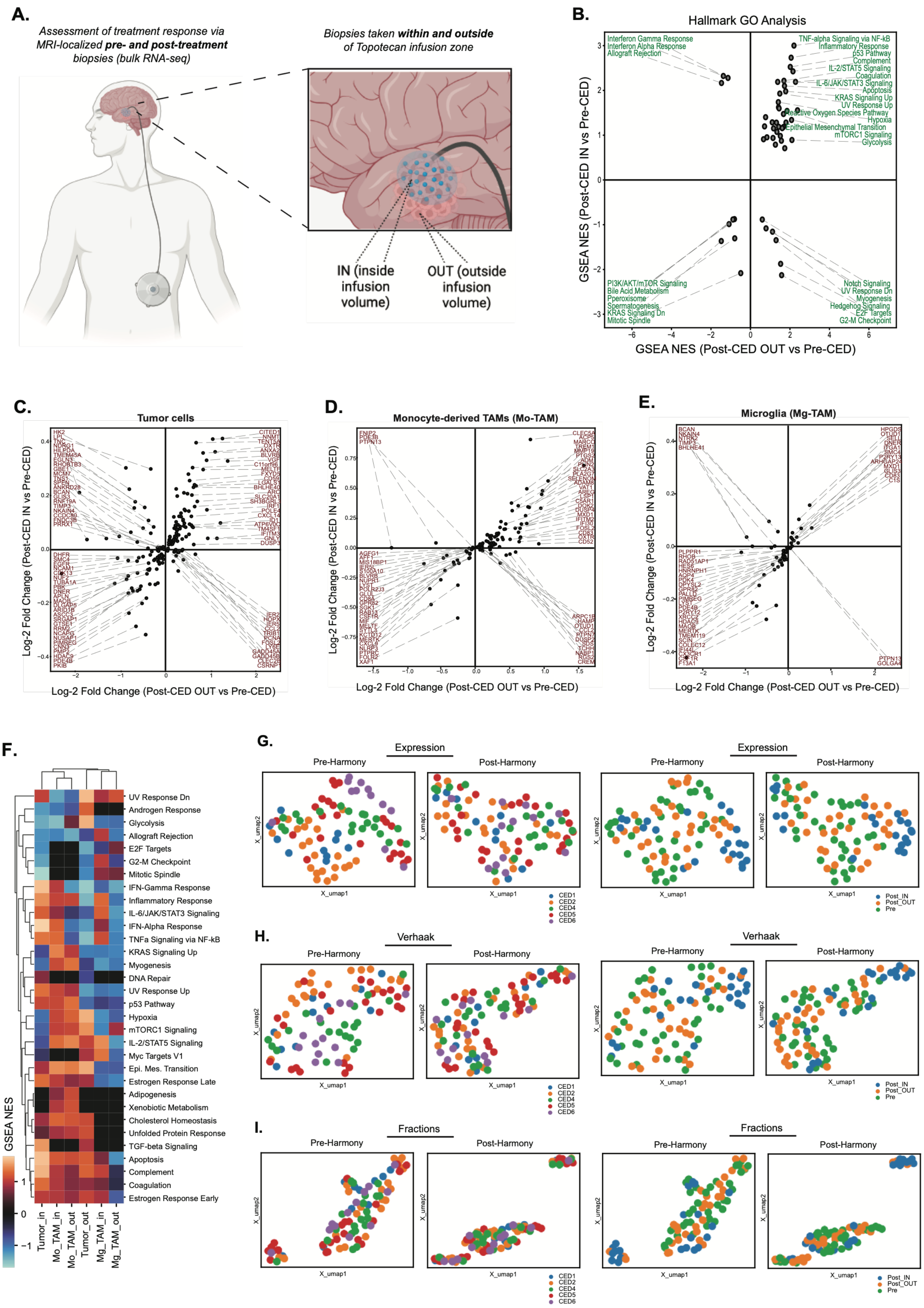
A.
Assessment of treatment response via MRI-localized pre- and post-treatment biopsies (bulk RNA-seq)
Biopsies taken within and outside of Topotecan infusion zone
IN (inside infusion volume)
OUT (outside infusion volume)
B.
Hallmark GO Analysis
Interferon Gamma Response
Interferon Alpha Response
Allograft Rejection
TNF-alpha Signaling via NF-kB
Inflammatory Response
p53 Pathway
Complement
IL-2/STAT5 Signaling
Coagulation
IL-6/JAK/STAT3 Signaling
Apoptosis
KRAS Signaling Up
UV Response Up
Reactive Oxygen Species Pathway
Hypoxia
Epithelial Mesenchymal Transition
mTORC1 Signaling
Glycolysis
PI3K/AKT/mTOR Signaling
Bile Acid Metabolism
Pperoxisome
Spermatogenesis
KRAS Signaling Dn
Mitotic Spindle
Notch Signaling
UV Response Dn
Myogenesis
Hedgehog Signaling
E2F Targets
G2-M Checkpoint
GSEA NES (Post-CED IN vs Pre-CED)
GSEA NES (Post-CED OUT vs Pre-CED)
C.
Tumor cells
D.
Monocyte-derived TAMs (Mo-TAM)
E.
Microglia (Mg-TAM)
Log-2 Fold Change (Post-CED IN vs Pre-CED)
Log-2 Fold Change (Post-CED OUT vs Pre-CED)
F.
UV Response Dn
Androgen Response
Glycolysis
Allograft Rejection
E2F Targets
G2-M Checkpoint
Mitotic Spindle
IFN-Gamma Response
Inflammatory Response
IL-6/JAK/STAT3 Signaling
IFN-Alpha Response
TNFa Signaling via NF-kB
KRAS Signaling Up
Myogenesis
DNA Repair
UV Response Up
p53 Pathway
Hypoxia
mTORC1 Signaling
IL-2/STAT5 Signaling
Myc Targets V1
Epi. Mes. Transition
Estrogen Response Late
Adipogenesis
Xenobiotic Metabolism
Cholesterol Homeostasis
Unfolded Protein Response
TGF-beta Signaling
Apoptosis
Complement
Coagulation
Estrogen Response Early
GSEA NES
Tumor_in
Mo_TAM_in
Mo_TAM_out
Tumor_out
Mg_TAM_in
Mg_TAM_out
G.
Expression
H.
Verhaak
I.
Fractions
Pre-Harmony
Post-Harmony
X_umap1
X_umap2
CED1
CED2
CED4
CED5
CED6
Post_IN
Post_OUT
Pre

**Figure S1. (A)** Study design and spatial biopsy acquisition from patients with recurrent glioblastoma treated with convection-enhanced delivery (CED) of topotecan, including pre-CED biopsies obtained before infusion and post-CED biopsies stratified by location within (post-CED IN) or outside (post-CED OUT) the maximal MRI-defined infusion zone. **(B)** Gene set enrichment analysis of Hallmark pathways showing upregulation of interferon responses, inflammatory signaling, hypoxia, and epithelial–mesenchymal transition together with suppression of cell-cycle programs in post-CED IN compared with pre-CED or post-CED OUT. **(C-E)** Cell-type–resolved transcriptional signatures showing induction of remodeling-associated macrophage genes, including MARCO and inflammatory mediators, together with loss of homeostatic microglial markers after treatment. **(F)** Heatmap of Hallmark pathway enrichment across inferred cell-type-specific differential expression profiles. **(G-I)** Sensitivity UMAP analysis showing effects of patient identity regression using multiple models. UMAPs show pre- and post-Harmony regression for bulk expression (G), Verhaak signatures (H), and CIBERSORTx-inferred fractions (I).

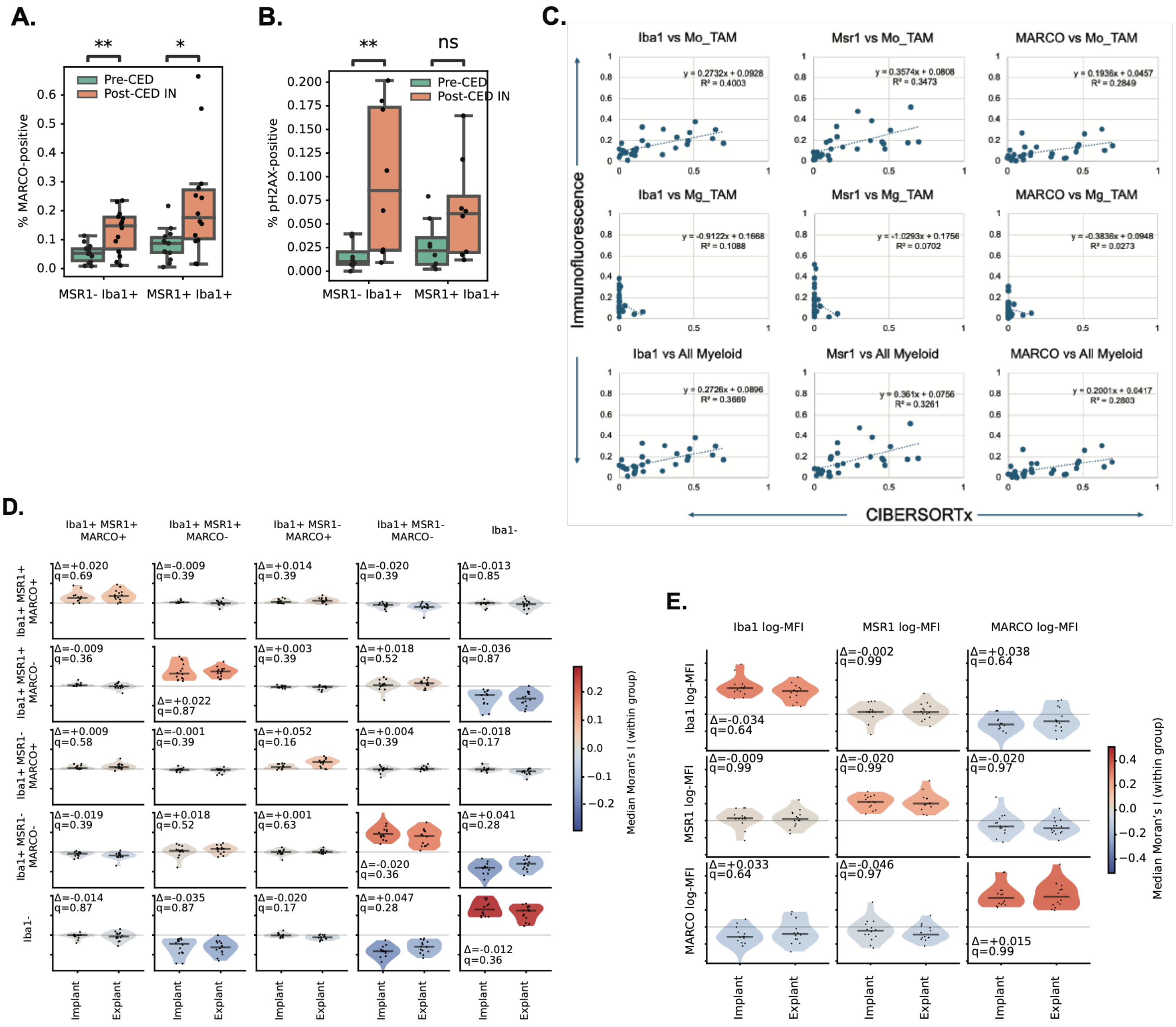


**Figure S2. (A–B)** Quantification of the percentage of MARCO+ cells (A) and pH2AX+ cells (B) among MSR1−IBA1+ and MSR1+IBA1+ myeloid populations in pre-CED and post-CED-IN biopsies, demonstrating broad MARCO induction across MSR1-defined myeloid compartments and preferential pH2AX enrichment within MSR1−IBA1+ myeloid cells following CED-topotecan treatment. **(C)** Correlation plots comparing CIBERSORTx-inferred cell-type abundance estimates from bulk RNA-seq with multiplex immunofluorescence–based cell quantification. Linear regression equations and $R^2$ values are shown, supporting concordance between transcriptomic deconvolution and orthogonal immunofluorescence measurements. **(D-E)** Spatial cross-correlation (Moran's I) of cell type identity based on binary marker positivity (D) and marker mean fluorescence (E) at the per-cell level, calculated for each slide and stratified by treatment condition.

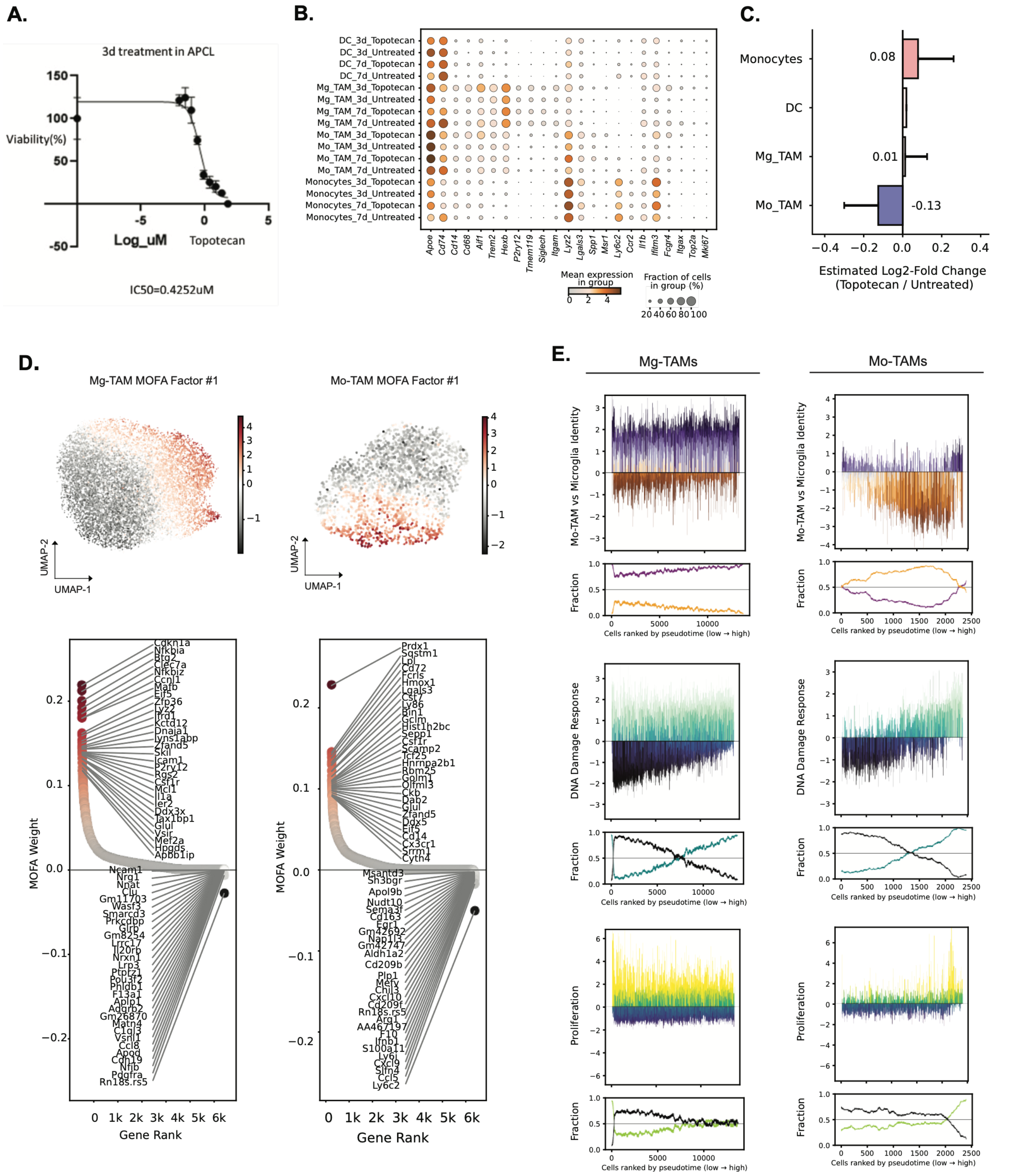


**Figure S3. (A)** Dose-response curve in murine p53-deficient, PDGFA-driven glioma cells demonstrating topotecan sensitivity, with an IC50 of 0.43 μM. **(B)** Dotplot depicting mean expression of canonical myeloid markers within each myeloid subset for individual mice. **(C)** Differential abundance of the myeloid compartment showing reduction of tumor cells and stability of the myeloid compartment after treatment. (**D)** Multi-omic factorization analysis (MOFA) of the Mo-TAM + Mg-TAM combined population and the top factor by variance explained, with scores projected onto UMAP embeddings and gene loadings depicting a homeostasis vs MES-inflammatory axis. **(E)** Barcode enrichment plots (top) illustrating cell-level representation of DNA damage repair, proliferation, and Mo-TAM vs Mg-TAM identity scores across cells ranked by pseudo-time (x-axis), with running fractional distributions of total signature-positive and signature-negative cells along pseudo-time trajectory shown on the bottom of each plot.

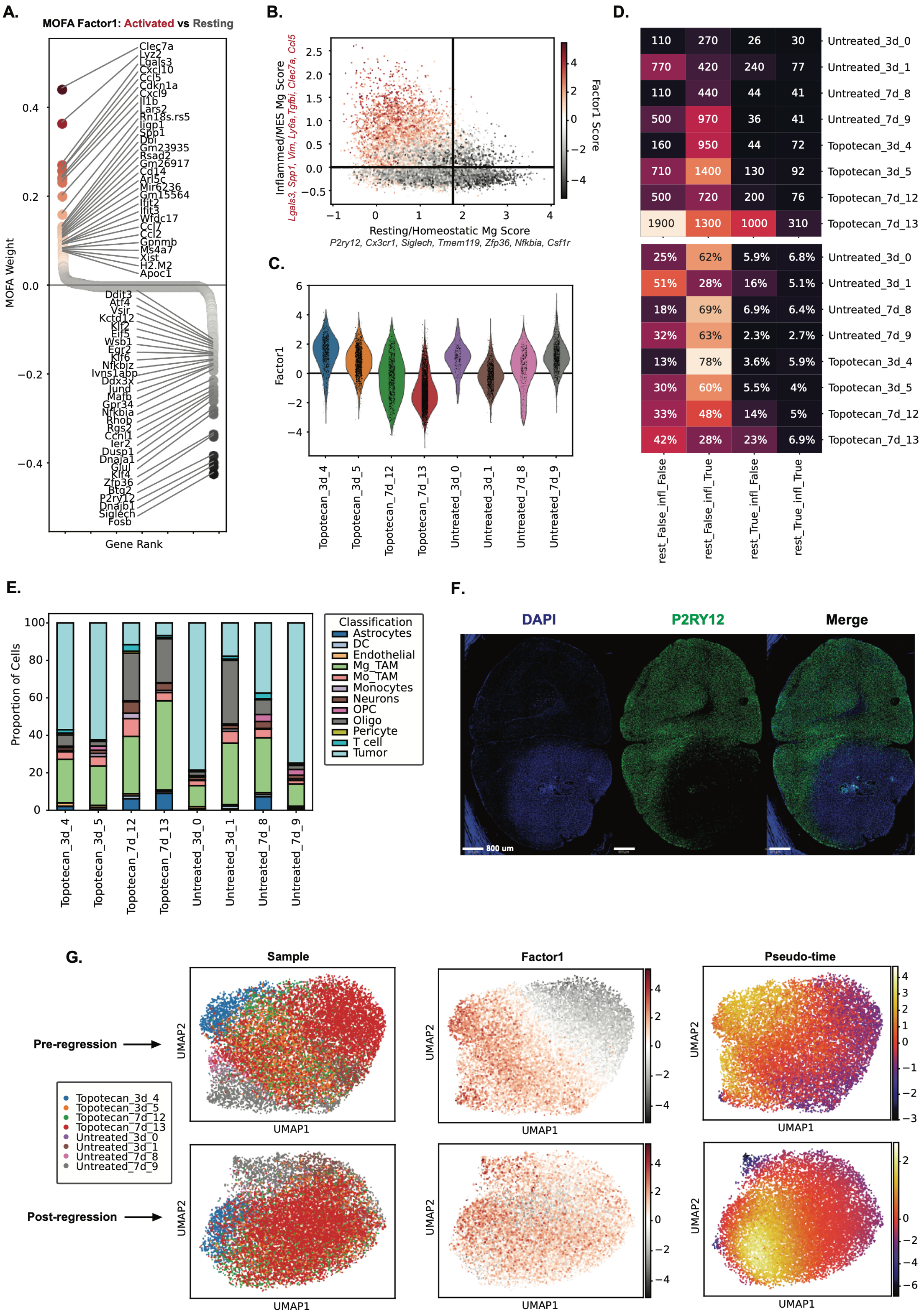
A.
MOFA Factor1: Activated vs Resting
MOFA Weight
Gene Rank
Clec7a
Lyz2
Lgals3
Cxcl10
Ccl5
Cdkn1a
Cxcl9
Il1b
Lars2
Rn18s.rs5
Iigp1
Spp1
Dbi
Gm23935
Rsad2
Gm26917
Cd14
Arl5c
Mir6236
Gm15564
Ifit2
Ifit3
Wfdc17
Ccl7
Ccl2
Gpnmb
Ms4a7
Xist
H2.M2
Apoc1
Ddit3
Atf4
Vsir
Kctd12
Klf2
Elf5
Wsb1
Egr2
Klf6
Nfkbiz
Ivns1abp
Ddx3x
Jund
Mafb
Gpr34
Nfkbia
Rhob
Rgs2
Ccnl1
Ier2
Dusp1
Dnaja1
Glul
Klf4
Zfp36
Btg2
P2ry12
Dnajb1
Siglech
Fosb
B.
Inflammed/MES Mg Score
Lgals3, Spp1, Vim, Ly6a, Tgfbi, Clec7a, Ccl5
Resting/Homeostatic Mg Score
P2ry12, Cx3cr1, Siglech, Tmem119, Zfp36, Nfkbia, Csf1r
Factor1 Score
C.
Factor1
Topotecan_3d_4
Topotecan_3d_5
Topotecan_7d_12
Topotecan_7d_13
Untreated_3d_0
Untreated_3d_1
Untreated_7d_8
Untreated_7d_9
D.
110 270 26 30 Untreated_3d_0
770 420 240 77 Untreated_3d_1
110 440 44 41 Untreated_7d_8
500 970 36 41 Untreated_7d_9
160 950 44 72 Topotecan_3d_4
710 1400 130 92 Topotecan_3d_5
500 720 200 76 Topotecan_7d_12
1900 1300 1000 310 Topotecan_7d_13
25% 62% 5.9% 6.8% Untreated_3d_0
51% 28% 16% 5.1% Untreated_3d_1
18% 69% 6.9% 6.4% Untreated_7d_8
32% 63% 2.3% 2.7% Untreated_7d_9
13% 78% 3.6% 5.9% Topotecan_3d_4
30% 60% 5.5% 4% Topotecan_3d_5
33% 48% 14% 5% Topotecan_7d_12
42% 28% 23% 6.9% Topotecan_7d_13
rest_False_infl_False
rest_False_infl_True
rest_True_infl_False
rest_True_infl_True
E.
Proportion of Cells
Classification
Astrocytes
DC
Endothelial
Mg_TAM
Mo_TAM
Monocytes
Neurons
OPC
Oligo
Pericyte
T cell
Tumor
F.
DAPI
P2RY12
Merge
800 um
G.
Sample
Factor1
Pseudo-time
Pre-regression
Post-regression
UMAP1
UMAP2

**Figure S4.** **(A)** Second highest variance-explained MOFA factor (Factor1) in Mg-TAMs, representing a resting vs activated Mg-TAM state. **(B)** Mg-TAMs were scored for inflammatory (y-axis) and resting (x-axis) genes and plotted in score-coordinate space. Single-cell MOFA Factor1 scores are shown as dot colors. **(C)** Violin plot depicting Factor1 score distributions for single Mg-TAM cells in each mouse. **(D)** Total cell count (top) and proportion (bottom) of cells within each quadrant in the plot in (B) for each mouse tumor sample. **(E)** Proportional abundance of each cell type in each mouse tumor profiled by scRNA-seq, depicting three mice (both Topotecan-7d and Un-3d_1) with over-sampled peripheral normal brain (high oligodendrocyte, neuron, and Mg-TAM proportions). **(F)** Representative immunofluorescence image demonstrating abundance of P2RY12+ resting Mg-TAMs in the periphery and absence of these cells in the tumor core. **(G)** Mg-TAM UMAP embeddings depicting sample distribution (left), Factor1 scores (middle), and single-cell pseudo-time scores (right), before (top row) and after (bottom row) regression of Factor1 scores.

### SUPPLEMENTAL METHODS: Paired Tumor Biopsies Reveal Spatiotemporal Myeloid Remodeling After Local Chemotherapy in Glioblastoma

#### SECTION A – EXPERIMENTAL PROCEDURES

##### 1. Human CED-topotecan biopsy specimens and multiplex immunofluorescence

*1.1 Patient samples and clinical biopsy annotation*

Recurrent glioblastoma specimens were obtained from patients enrolled in a previously reported investigator-initiated, single-center phase 1b clinical trial of convection-enhanced delivery of topotecan conducted at NewYork-Presbyterian/Columbia University Irving Medical Center (1). Pre-CED biopsies were obtained at the time of catheter implantation, and post-CED biopsies were obtained at catheter explantation after completion of topotecan infusion. Paired pre-CED and post-CED biopsies from five patients were analyzed. Post-CED biopsies were annotated as within or outside the maximal infusion zone, designated post-CED IN and post-CED OUT, based on intraoperative biopsy coordinates and post-infusion MRI localization of co-infused gadolinium, as previously described. Biopsies were analyzed by bulk RNA sequencing and immunofluorescence as described below.

The parent clinical trial enrolled adults aged 18 years or older with previously confirmed WHO grade III–IV malignant glioma who had undergone prior surgery, temozolomide, and radiotherapy and subsequently presented with clinical and radiographic recurrence requiring reoperation. Eligible tumors were solitary, supratentorial, contrast-enhancing lesions less than 32 $cm^3$ and were required to be stereotactically accessible. Additional eligibility criteria included Karnofsky Performance Status ≥70, adequate hematologic, metabolic, and coagulation parameters, and eligibility for MRI and PET imaging. Key exclusion criteria included prior systemic topotecan therapy, multifocal, cerebellar, or ventricular disease, active infection including HIV or hepatitis B/C, and concurrent use of narrow-therapeutic-index drugs metabolized exclusively by CYP450 enzymes.

All patients provided written informed consent under a protocol approved by the Columbia University Institutional Review Board (protocol #AAAQ9520) and the FDA. The principal investigator held the investigational new drug sponsorship, and the clinical trial was registered as NCT03154996.

*1.2 Multiplex immunofluorescence staining*

Paraffin-embedded brain sections were cut at 5 µm, mounted onto slides, deparaffinized, and rehydrated through graded ethanol. Heat-induced epitope retrieval was performed using 10 mM citrate buffer at pH 6.0, unless otherwise indicated below. Sections were blocked in 10% normal goat serum in PBS and incubated with primary antibodies, followed by species-appropriate fluorescent secondary antibodies. Nuclei were counterstained with DAPI. Tyramide signal amplification was used when sequential staining of same-host primary antibodies was required. In these cases, the first primary antibody was detected using a tyramide-based amplification kit, followed by signal development and appropriate antibody inactivation/blocking before application of subsequent primary antibodies.

Stained slides were imaged on a Nikon AX confocal microscope with z-stack acquisition. Imaging parameters, including laser power, detector gain, pinhole size, z-step interval, and exposure settings, were kept constant across slides stained within the same antibody panel. Images were quantified in QuPath as described below. Regions affected by large artifacts, tissue folding, tissue edges, hemorrhage, or nonrepresentative necrotic debris were excluded from QuPath annotation.

CED-topotecan biopsy samples obtained from pre-CED catheter implantation or post-CED infusion-zone tissue were stained for pH2AX, MARCO, MSR1, and Iba1. Primary antibodies included pH2AX (1:1000, Cell Signaling Technology, 9718S), MARCO (1:1000, rabbit anti-MARCO, Invitrogen, PA5-64134; antigen retrieval pH 9), Iba1 (1:1000, rabbit anti-Iba1, Cell Signaling Technology, 17198), Iba1 (1:1000, Aves, IBA1-0200), and MSR1 (1:1000, mouse IgG1 anti-MSR1, Thermo Fisher, J5HTR3). For MARCO/Iba1 staining, MARCO was applied first and detected using the Alexa Fluor™ 488 Tyramide SuperBoost™ Kit with goat anti-rabbit IgG amplification (Invitrogen, B40922), followed by staining for Iba1 and the remaining primary antibodies. Secondary antibodies included goat anti-rabbit Alexa Fluor 488 (Invitrogen, A11008), goat anti-mouse IgG1 Alexa Fluor 555 (Invitrogen, A21127), and goat anti-chicken Alexa Fluor 647 (Invitrogen, A21449). Whole-biopsy regions were analyzed, and quantification quality was assessed by independent blinded reviewers through review of individual segmentation and classification results.

*1.3 QuPath-derived cell counts and labeling indices*

Multiplex immunofluorescence quantification was performed from QuPath-derived detection tables containing slide-level area measurements and counts of total valid detections, single-marker-positive cells, and marker-overlap-positive cells.

Cell detection was performed in QuPath using StarDist-based nuclear segmentation on the DAPI channel. A pretrained StarDist model was applied to identify nuclei, followed by cell expansion to approximate whole-cell boundaries for marker quantification (2). StarDist and cell-expansion parameters were optimized on representative images and then held constant across all images within a staining panel. Marker-positivity thresholds were defined using negative-control regions and visual review of representative positive and negative cells and were applied uniformly across comparable slides. Classification outputs were reviewed for segmentation accuracy and marker-threshold performance before export.

Patient and biopsy identifiers were concatenated to generate patient-biopsy identifiers for downstream mixed modeling. Overall cellularity was calculated as 1,000 multiplied by the number of valid cell detections divided by annotated tissue area, yielding events per standardized tissue area. Labeling indices for Iba1, MSR1, MARCO, and pH2AX were calculated as 100 times the number of marker-positive cells divided by the number of valid detections.

Marker positivity within the myeloid compartment was quantified by dividing the number of Iba1-positive cells co-positive for the marker of interest by the total number of Iba1-positive cells. MSR1-positive Iba1-positive, MARCO-positive Iba1-positive, and pH2AX-positive Iba1-positive labeling indices were calculated in this manner. For subset analyses stratifying myeloid cells by MSR1 status, MARCO or pH2AX positivity was calculated separately among MSR1-positive and MSR1-negative Iba1-positive cells using the corresponding overlap count tables.

##### *1.4 Statistical modeling of immunofluorescence readouts*

Immunofluorescence endpoints were tested using a hierarchical model-fitting strategy designed to account for repeated measurements while maintaining robustness when more complex models failed to converge. For each marker or labeling-index endpoint, the primary model was a mixed-effects model with condition as a fixed effect, patient as a random intercept, and biopsy-level variance components when the data supported that structure. If this full model failed to converge or could not be fit for a particular endpoint, a simpler patient-random-intercept mixed model was attempted. If mixed modeling remained unstable, ordinary least-squares regression with cluster-robust standard errors by patient was used as the fallback. Pairwise condition contrasts were extracted from model coefficients and covariance matrices and adjusted using Holm correction. Omnibus tests were derived from likelihood-ratio comparisons for mixed models where possible.

Boxplots and stripplots display the distribution of analysis-level measurements by condition, with p-value brackets generated from the corresponding Holm-adjusted pairwise contrasts. This strategy was used for the human biopsy immunofluorescence analyses and adapted for mouse immunofluorescence endpoints where treatment groups rather than patient-paired clinical regions defined the primary comparison.

##### *1.5 Spatial and fluorescence-intensity secondary analyses*

Supplemental QuPath analyses were performed to summarize spatial relationships and fluorescence-intensity patterns among marker-defined cell classes. Cell classifications exported from QuPath were recoded into interpretable marker-combination categories, including Iba1-positive/MSR1-positive/MARCO-negative cells, Iba1-positive/MSR1-negative/MARCO-negative cells, Iba1-positive/MSR1-positive/MARCO-positive cells, and Iba1-positive/MSR1-negative/MARCO-positive cells. QuPath-derived class-by-class spatial summary matrices, marker mean-fluorescence intensity matrices, and associated p-value matrices were imported for pre-CED and post-CED samples. Spatial co-localization patterns and marker fluorescence-intensity relationships among marker-defined cell classes were visualized as clustered heatmaps. Significance overlays were derived from the associated p-value matrices, allowing the supplemental figure panels to display both the direction and strength of spatial and fluorescence-intensity relationships across treatment conditions.

#### 2. Murine CED-topotecan, survival, immunofluorescence, and single-cell RNA-seq

##### *2.1 Animal ethics and murine model*

All animal experiments were performed in accordance with institutional and national guidelines for the care and use of laboratory animals and were approved by the Columbia University Institutional Animal Care and Use Committee (Protocol #AC-AABO9554). Female C57BL/6 mice aged 6–8 weeks were used for all experiments. Orthotopic gliomas were established using a PDGFA-driven, p53-deficient syngeneic glioma model, as previously described (3). Briefly, mice were anesthetized with ketamine/xylazine (100 mg/kg and 10 mg/kg, respectively) and assessed for lack of reflexes by toe pinch. Hair was shaved, and a vertical scalp incision was performed. A burr hole was made with a 17-gauge needle 2 mm lateral and 2 mm anterior to the bregma.

A cell suspension was prepared from cultured APCL cells. Tumor cells were injected stereotactically with a Hamilton syringe at 0.3 μL/min to deliver 50,000 APCL cells in <2 μL, 2 mm deep into subcortical white matter. Tumor growth was assessed longitudinally by luciferase bioluminescence imaging, as previously described (4).

*2.2 Convection-enhanced delivery of topotecan in mice*

Pump implantation was performed as previously described (5, 6). Alzet osmotic pumps (1007D) were filled with vehicle control or topotecan diluted to 146 μM in PBS. Topotecan was prepared from a 100 mM stock dissolved in 100% DMSO and diluted to the final working concentration before pump loading; the vehicle control contained the matched final DMSO concentration. Pumps were loaded under sterile conditions, connected to brain-infusion cannulas, and primed overnight in sterile PBS at 37°C before implantation. At 21 days after tumor injection, osmotic pumps were implanted subcutaneously, and the catheter was placed through the same burr hole used for tumor implantation.

The catheter was secured using instant adhesive (Loctite 454, Alzet #0008670). The incision was closed with sutures. Pumps delivered vehicle or topotecan continuously for seven days.

One percent Omniscan was included in all Alzet pumps to confirm intraparenchymal distribution by MRI.

*2.3 Murine MRI scans*

MRI scans were conducted using a Bruker BioSpec 9.4T scanner. Mice were anesthetized with 1–2% isoflurane mixed with medical air and administered via a nose cone. Isoflurane concentration was adjusted throughout imaging to maintain a stable respiratory rate between 40 and 70 breaths per minute. Respiration was monitored using a sensor pillow connected to a physiological monitoring system, and body temperature was maintained at approximately 37°C using a circulating water heating pad.

Low-resolution T1-weighted scout images were acquired for initial localization. High-resolution anatomical imaging was performed using a T2-weighted rapid acquisition with relaxation enhancement sequence with the following acquisition parameters: repetition time = 3000 ms, echo time = 45 ms, field of view = 17 × 15 mm, matrix resolution = 225 × 198, slice thickness = 0.7 mm, and 16 slices spanning the entire brain. Contrast-enhanced imaging was performed using a T1-weighted sequence with the same geometric parameters and the following acquisition settings: repetition time = 150 ms, echo time = 2.2 ms, and flip angle = 70°.

*2.4 Murine survival analysis*

Murine survival analyses were performed from the time of tumor implantation until death or predefined humane endpoint. Mice were checked daily for signs of neurologic decline or tumor-related morbidity. Humane endpoints included decreased activity or alertness, hunched posture, seizures, or inability to feed secondary to motor dysfunction. The survival results shown represent the combination of two survival studies, including 14 mice in the control group and 13 mice in the topotecan group. Animals receiving CED-topotecan or vehicle control were plotted using Kaplan–Meier survival curves, and survival distributions were compared using the log-rank test. Median survival values were reported for each treatment group.

*2.5 Tissue collection and processing*

For post-treatment tissue analyses, mice were euthanized immediately following completion of seven days of CED treatment. Animals were anesthetized and perfused transcardially with 15 mL PBS followed by 15 mL 4% paraformaldehyde. Brains were extracted, further fixed in 4% paraformaldehyde for 24 hours at 4°C, processed for paraffin embedding, and sectioned at 5 μm. Sections were selected at levels containing the catheter tract and adjacent tumor-bearing tissue for immunohistochemistry and immunofluorescence.

For single-cell RNA-seq, animals were euthanized following seven days of CED-topotecan or vehicle treatment without fixation/perfusion with paraformaldehyde. Tumor-bearing brain regions were rapidly dissected and immediately processed into single-cell suspensions as described below.

*2.6 Murine immunofluorescence and histological quantification*

Mouse brain sections were stained for pH2AX, MSR1, and Iba1. For mouse sections, blocking was performed in 10% normal goat serum in PBS containing 0.3% Triton X-100, and primary antibodies were diluted in the same blocking solution. Primary antibodies included pH2AX (1:1000, Cell Signaling Technology, 9718S), Iba1 (1:1000, Aves, IBA1-0200), and MSR1 (1:2000, Cell Signaling Technology, 98215S). For pH2AX/MSR1 co-staining, MSR1 was applied first and detected using the Alexa Fluor™ 594 Tyramide SuperBoost™ Kit with goat anti-rabbit IgG amplification (Invitrogen, B40944), followed by staining for pH2AX and the remaining primary antibodies. Secondary antibodies included goat anti-rabbit Alexa Fluor 488/568 (Invitrogen, A11008/A11036) and goat anti-chicken Alexa Fluor 647 (Invitrogen, A21449).

Slides were imaged using identical acquisition settings across treatment groups within each staining batch. Quantitative histologic analysis of murine CED tumors was performed in QuPath within standardized regions of interest centered on the catheter tract to capture the local treatment-affected microenvironment. Regions of interest were drawn blinded to treatment group and kept consistent in size and anatomical orientation across animals. Regions affected by large artifacts, tissue folding, tissue edges, hemorrhage, or nonrepresentative necrotic debris were excluded from QuPath annotation. Cell detection was performed using StarDist-based nuclear segmentation on the DAPI channel, followed by cell expansion to approximate whole-cell boundaries for marker quantification (2). StarDist and classification parameters were optimized on representative images and held constant across treatment groups within each staining batch. Marker-positivity thresholds for Iba1, MSR1, and pH2AX were established by visual review of representative positive and negative cells.  Total cellularity was calculated as segmented events per tissue area. Iba1, MSR1, and pH2AX labeling indices were calculated as the fraction of segmented cells positive for each marker. To evaluate marker expression within the myeloid compartment, Iba1-positive cells were used as the myeloid denominator. Myeloid-restricted MSR1 and pH2AX positivity were calculated by dividing the number of Iba1/MSR1-positive or Iba1/pH2AX-positive events by the total number of Iba1-positive events. Group-level comparisons were performed at the animal level, rather than treating individual segmented cells as independent biological replicates.

##### *2.7 Mouse single-cell RNA-seq sample processing*

For single-cell RNA-seq, mice were euthanized following three or seven days of CED-topotecan or vehicle treatment. Mouse brains were rapidly dissected, and the tumor-bearing quadrant of the brain was isolated. Tissue was enzymatically dissociated to generate a single-cell suspension, and viable cells were enriched before library preparation. Libraries were generated using a droplet-based single-cell platform and sequenced according to the manufacturer's protocol.

#### 3. Ex vivo slice culture and in vitro perturbation

##### *3.1 Patient-derived ex vivo slice culture and drug treatment*

For the present study, paired DMSO- and topotecan-treated slice-culture scRNA-seq datasets were generated from three newly processed patient-derived glioblastoma specimens and integrated with two previously published paired slice-culture datasets generated using the same experimental framework. Newly processed glioblastoma specimens were obtained from surgical resections performed at Columbia University Irving Medical Center under a protocol approved by the Columbia University Irving Medical Center Institutional Review Board. Tumor specimens were collected immediately after surgical removal and transported in ice-cold artificial cerebrospinal fluid containing 210 mM sucrose, 10 mM glucose, 2.5 mM KCl, 1.25 mM $NaH_2PO_4$, 0.5 mM $CaCl_2$, 7 mM $MgCl_2$, and 26 mM $NaHCO_3$.

Ex vivo tissue-slice preparation was performed using a previously described protocol (7). Briefly, tumor specimens were placed in ice-cold artificial cerebrospinal fluid and sectioned under sterile conditions into 500 μm slices using a McIlwain tissue chopper. Slices were transferred to ice-cold artificial cerebrospinal fluid in 6-well plates using a sterile plastic Pasteur pipette and allowed to recover for 15 minutes as the solution reached room temperature. Intact, well-shaped slices approximately 5–10 mm in diameter were then transferred onto porous membrane inserts, 0.4 μm pore size, in 6-well plates containing 1.5 mL maintenance medium consisting of DMEM/F12 supplemented with N-2 supplement and 1% antibiotic-antimycotic. Culture medium was placed beneath the membrane insert without bubbles to ensure diffusion into the slice, and 10 μL of culture medium was added directly onto each slice to prevent drying.

Slices were rested for 6 hours in maintenance medium in a humidified incubator at 37°C and 5% $CO_2$. The medium was then replaced with pre-warmed medium containing 20 μM topotecan or matched DMSO vehicle control. Slices were cultured in treatment medium for 18 hours at 37°C and 5% $CO_2$ before collection and dissociation for downstream single-cell RNA-seq analysis.

##### *3.2 iPSC-derived microglia differentiation and treatment*

Induced pluripotent stem cells were differentiated into primitive hematopoietic progenitor cells using the STEMdiff™ Hematopoietic Kit (STEMCELL Technologies, 05310), adapted from McQuade et al (8). Hematopoietic progenitor cells were plated on Cultrex-coated 24-well plates at a density of 100,000 cells per well and maintained at 37°C in DMEM/F12-based microglial differentiation medium supplemented with insulin-transferrin-selenium, B27, N2, GlutaMAX, MEM non-essential amino acids, monothioglycerol, insulin, and penicillin-streptomycin. Cells were fed every two days with a tri-cytokine differentiation cocktail containing 100 ng/mL IL-34, 50 ng/mL TGF-β1, and 25 ng/mL M-CSF. On day 13, cells were gently lifted by mechanical disruption and split as needed to maintain appropriate confluency. Differentiation continued with the tri-cytokine cocktail until day 25, after which 100 ng/mL CD200 and 100 ng/mL CX3CL1 were added to promote microglial maturation. Cells were fed daily until day 28, when mature iPSC-derived microglia were used for transcriptomic assays. For perturbation experiments, mature iPSC-derived microglia were treated with 2.5 μM topotecan or matched DMSO vehicle control for 2 days before RNA extraction and bulk RNA-seq. In parallel, iPSC-derived microglia were treated with lipopolysaccharide (LPS; 1 ng/μL) as a positive control for canonical inflammatory activation.

*3.3 In vitro dose-response and viability*

Murine p53-deficient, PDGFA-driven glioma cells were maintained as previously described (3). Cells were plated in triplicate in clear flat-bottom 96-well plates at 5,000 cells/well. Twenty-four hours after plating, cells were treated with DMSO vehicle or serial concentrations of topotecan. Viability was assessed using the CellTiter 96® AQueous One Solution MTS Assay (Promega, G3582). Absorbance values were normalized to the mean DMSO vehicle-control signal. Dose-response curves were generated by plotting normalized viability as a function of topotecan concentration, and the IC50 was estimated from the fitted dose-response curve.

Human HMC3 microglia and U87 glioma cells were cultured under standard conditions in DMEM with 10% fetal bovine serum (7, 8). HMC3 microglia, U87 glioma cells, or HMC3–U87 co-cultures were plated in triplicate in white flat-bottom 96-well plates at 5,000 total cells/well, with co-cultures plated at a 1:1 HMC3:U87 ratio. Twenty-four hours after plating, cells were treated with DMSO vehicle or serial concentrations of topotecan ranging from 10 nM to 10 μM for 72 hours. Viability was assessed using the CellTiter-Glo Luminescent Cell Viability Assay (Promega, G7570). Luminescence was measured on a Tecan Infinite 200 Pro plate reader using 100 ms integration time. Raw luminescence values were normalized to the mean DMSO vehicle-control signal for each plate.

Dose-response curves in HMC3 microglial cells, U87 glioma cells, and HMC3–U87 co-culture were generated by plotting normalized viability as a function of topotecan concentration, and replicate-level means and standard deviations were calculated for each dose. These dose-response data were used to identify a topotecan concentration that produced glioma-cell effects while preserving sufficient microglial viability for downstream inflammatory assays.

*3.4 In vitro cytokine production*

For cytokine assays, cells were plated in monoculture or HMC3–U87 co-culture in 6-well plates at a density of $3 \times 10^5$ cells per well and treated for 72 hours with vehicle, LPS (1 ng/μL), or topotecan (100 nM). Each condition was performed with 3–4 biological replicates. Following treatment, RNA was extracted from cell lysates using the RNeasy Mini Kit (Qiagen, 74106). cDNA was synthesized using the Invitrogen SuperScript VILO cDNA Synthesis Kit according to the manufacturer's protocol. Quantitative PCR was performed on a QuantStudio 6 Pro Real-Time PCR System using Thermo Scientific ABsolute Blue qPCR Mix, SYBR Green, Low ROX (AB4163A), with ACTB as the reference gene. Primer sequences were as follows: IL1B forward, ATGATGGCTTATTACAGTGGCAA, reverse, GTCGGAGATTCGTAGCTGGA; IL6 forward, ACTCACCTCTTCAGAACGAATTG, reverse, CCATCTTTGGAAGGTTCAGGTTG; and ACTB forward, TGGCACCCAGCACAATGAA, reverse, CTAAGTCATAGTCCGCCTAGAAGCA.

IL1B and IL6 transcript expression was quantified by the $\Delta\Delta Ct$ method, with fold change calculated as $2^{-\Delta\Delta Ct}$ relative to the control group mean. Statistical analyses of qPCR data were performed on $\Delta Ct$ values. For visualization, expression values were summarized relative to control-treated cells, such that higher plotted values represented induction relative to control. Topotecan- and LPS-treated groups were compared within each target gene using two-sided Welch t-tests.

IL1B and IL6 protein levels were quantified from co-culture lysates using a Meso Scale Discovery electrochemiluminescence immunoassay platform according to the manufacturer's protocol. Plates were read on an MSD QuickPlex instrument, and cytokine concentrations were calculated from standard curves generated on each plate and normalized to total protein. Replicate-level cytokine abundance was summarized for each condition. For MSD cytokine measurements, control, topotecan, and LPS groups were compared within each target using one-way analysis of variance. When the omnibus test was significant, pairwise Welch t-tests were performed with Bonferroni correction for multiple comparisons.

*3.5 In vitro macrophage phagocytosis assay*

For the macrophage phagocytosis assay, APCL tumor cells were lifted with TrypLE, resuspended in complete media, counted, and labeled with CFSE at a final concentration of 10 µM for 3 hours at 37°C protected from light. CFSE-labeled APCL tumor cells were washed, resuspended in complete media, and plated in 12-well plates at $4.0 \times 10^5$ cells/well. Tumor cells were allowed to adhere for 18 hours, treated with 5 µM topotecan for 72 hours, washed with PBS, and then co-cultured with RAW 264.7 macrophages for 3 hours at 37°C.

Phagocytosis was analyzed by flow cytometry/FlowJo and calculated as the percentage of CFSE-positive cells within the eFluor-positive phagocyte population. After co-culture, cells were collected, washed, and analyzed by flow cytometry. Debris and doublets were excluded by forward- and side-scatter gating, viable single cells were retained when viability staining was performed, and macrophages were identified as eFluor-positive events. Gating thresholds were established using single-color and unstained controls and applied consistently across samples.

Macrophage phagocytosis assays were analyzed from replicate-level tables containing the fraction or percentage of macrophages positive for uptake of labeled tumor-cell material. Group comparisons were performed using the appropriate two-group or multi-group framework matching the plotted conditions, and replicate-level values were shown in the figure together with group summaries.

#### SECTION B – COMPUTATIONAL METHODS

##### 4. General computational framework and data conventions

*4.1 Computational environment and analysis objects*

All transcriptomic analyses were performed in Python using AnnData- and MuData-centered data structures. Count matrices were stored as observations-by-genes matrices, with raw count matrices retained in AnnData layers whenever downstream differential expression required integer counts. Processed metadata were stored in observation-level tables and were harmonized before analysis so that treatment condition, patient, biopsy, time point, sample, and inferred cell type annotations could be accessed consistently across workflows. Unless otherwise indicated, single-cell count matrices were normalized with scanpy.pp.normalize_total() to $1 \times 10^4$ counts per cell and transformed with scanpy.pp.log1p() for visualization, clustering, gene scoring, and label-transfer operations. Bulk RNA-seq expression matrices were normalized to $1 \times 10^6$ counts per sample and log-transformed for signature projection and visualization, while raw integer counts were used for DESeq2-style differential expression.

*4.2 Gene filtering and gene-set resources*

Gene filtering was performed to reduce technical signal from known confounding gene classes. Mitochondrial genes were identified by the prefix MT- or species-specific lower-case equivalents, ribosomal genes were identified using RPS/RPL or Rps/Rpl prefixes, hemoglobin genes were identified using regular expressions matching hemoglobin genes while excluding pseudogene-like forms where appropriate, and heat-shock genes were identified using HSP/Hsp prefixes. Non-coding-like genes with accession-style prefixes such as AL, AC, AF, AP, AJ, BX, Z8, and Z9 followed by numeric identifiers were excluded in the human workflows when these features were not biologically interpretable for downstream enrichment analyses. Gene filtering thresholds were defined separately for each analysis block and are described below.

Gene set enrichment analyses used preranked GSEA implemented with gseapy. Hallmark pathway libraries were retrieved from Enrichr/gseapy by preferentially selecting the most recent available MSigDB Hallmark library for the relevant organism, with MSigDB_Hallmark_2020 used as the fallback library name. For human analyses, the human Hallmark library was used directly. For mouse analyses, the mouse Hallmark library was used when available; where human pathway definitions were required for scoring mouse cells, human genes were mapped to mouse orthologs using gseapy Biomart-derived homolog tables and pathways with at least five mapped genes were retained. For preranked gene set enrichment analysis, genes were ranked by the log2 fold-change statistic from the relevant differential expression model. Enrichment analyses used 1,000 permutations, a fixed random seed of 42, and analysis-specific gene set size thresholds (minimum of 5) as described below. Normalized enrichment scores and Benjamini–Hochberg-adjusted false discovery rates were used for pathway-level visualization. Curated literature glioblastoma and immune signatures, including Verhaak bulk subtype signatures, Neftel malignant-state signatures, and manuscript-specific marker sets, were projected onto either single cells or bulk expression matrices using scanpy.tl.score_genes() (9, 10).

#### 5. Human CED-topotecan bulk RNA-seq analysis

##### *5.1 Bulk RNA-seq count preprocessing*

Raw bulk RNA-seq count matrices were loaded together with sample-level clinical metadata and converted into an AnnData object with biopsies as observations and genes as variables. Genes belonging to ribosomal, mitochondrial, hemoglobin, heat-shock, and non-coding-like categories were removed before downstream analyses as described above. Library-size normalization was then performed to $1 \times 10^6$ counts per sample, followed by log2(count + 1) transformation for visualization and signature scoring. This transformed matrix was used for subtype projection and exploratory visualization, whereas differential expression was performed on raw counts after gene filtering.

##### *5.2 Differential expression across pre-CED, post-CED IN, and post-CED OUT biopsies*

Differential expression was performed for the bulk transcriptome and for inferred Tumor, Mo_TAM, and Mg_TAM expression profiles derived from CIBERSORTx high-resolution deconvolution (see below). For the bulk analysis, the filtered count matrix was used directly. For inferred cell-type analyses, CIBERSORTx high-resolution expression profiles were read for each cell type, aligned to the same sample order as the clinical metadata, and rescaled from transcripts-per-million-like expression units back to integer count scale by multiplying each inferred expression value by the original library size of that bulk sample divided by 1,000,000. This produced an approximate count matrix that was used for negative-binomial modeling while preserving sample-specific sequencing depth.

Differential expression was performed with PyDESeq2 using raw integer counts. Genes with fewer than 100 total counts across all retained samples were excluded before model fitting. A single model matrix included patient identity and biopsy phenotype, thereby accounting for patient-level baseline differences when estimating treatment-region contrasts. DESeq2 dispersion estimation, model fitting, and Wald statistics were calculated using DefaultInference with eight CPUs and Cook's-distance refitting enabled. Three contrasts were extracted from the fitted model: post-CED IN versus pre-CED, post-CED OUT versus pre-CED, and post-CED IN versus post-CED OUT. Differential expression results were exported as gene-wise log2 fold changes, standard errors, test statistics, nominal p-values, and Benjamini–Hochberg-adjusted p-values. Genes with adjusted $p < 0.05$ were treated as statistically significant for gene-level visualization and for defining the gene universe plotted in spatial-region quadrant analyses.

*5.3 Visualization of region-specific transcriptional remodeling*

To visualize transcriptional remodeling associated with drug exposure, gene-wise log2 fold changes from post-CED IN versus pre-CED were plotted against corresponding log2 fold changes from post-CED OUT versus pre-CED. The plotted gene set was restricted to genes significant in at least one of these two contrasts, which emphasized genes with statistically supported changes in either post-treatment spatial compartment. Genes in the upper-right and lower-left quadrants represented concordant upregulation or downregulation in both post-CED regions relative to pre-CED, whereas genes in the upper-left and lower-right quadrants represented spatially discordant changes between the infusion zone and outside-zone specimens. Genes were labeled by magnitude and significance to highlight the most strongly region-associated transcriptional changes by Euclidean distance from the origin. Volcano plots were used for individual contrasts when only one differential expression signature was shown, with the x-axis representing log2 fold change and the y-axis representing −log10 p-value.

*5.4 Hallmark pathway enrichment of bulk differential expression signatures*

Hallmark pathway results were used to summarize inflammatory, interferon, hypoxia, cell-cycle, DNA-damage, and mesenchymal remodeling programs across bulk and cell-type-resolved expression profiles. Preranked GSEA was performed on each bulk and inferred cell-type differential-expression signature using log2 fold change as the ranking statistic as described above. For the clinical CED cohort, enrichment analyses were performed separately for Bulk, Tumor, Mg_TAM, and Mo_TAM signatures for both post-CED IN versus pre-CED and post-CED OUT versus pre-CED contrasts. For the bulk human biopsy comparisons, enrichment analyses were run with a minimum gene-set size of 10 and a maximum gene-set size of 5,000. Normalized enrichment scores were used to compare pathway direction and magnitude across contrasts, and Benjamini–Hochberg-adjusted false discovery rates were used to identify statistically enriched pathways. Literature signature enrichments (Verhaak, Neftel) were processed with the same preranked framework and combined with Hallmark results in a single enrichment output table.

*5.5 Projection of canonical bulk glioblastoma subtype signatures*

Canonical bulk glioblastoma subtype signatures (mesenchymal, classical, proneural, and neural) were projected onto the normalized and log-transformed bulk RNA-seq matrix using scanpy.tl.score_genes(). For each biopsy, the resulting signature score represented the average expression of genes in the subtype signature relative to a matched control gene set selected by Scanpy. All-biopsy patient-fixed-effects models were used to compare Verhaak signature scores across biopsy groups, using patient and biopsy phenotype as design terms (~ Patient + Phenotype) to account for repeated biopsies from the same patient. Planned contrasts included post-CED IN versus pre-CED, post-CED OUT versus pre-CED, and post-CED IN versus post-CED OUT. The post-CED IN versus pre-CED and post-CED OUT versus pre-CED contrasts tested treatment-associated transcriptional remodeling relative to pre-CED biopsies, whereas the direct post-CED IN versus post-CED OUT contrast tested spatial enrichment within the infusion zone. In parallel, patient-adjusted differential expression was performed with PyDESeq2 as described above, and the resulting log2 fold-change signatures were used for preranked GSEA of Verhaak subtype gene sets. For all analyses, FDR-adjusted q-values were used to assess significance, with $q < 0.05$ considered significant.

#### 6. CIBERSORTx digital cytometry and high-resolution expression inference

##### *6.1 Recurrent glioblastoma reference construction*

CIBERSORTx deconvolution was performed using a recurrent glioblastoma single-cell reference derived from the GBmap atlas and related recurrent glioblastoma annotations (11). The reference was restricted to recurrent biopsies to better match the clinical setting of the CED-topotecan cohort. Published and harmonized cell-type labels were mapped into the cell populations used for deconvolution, including tumor cells, Mo-TAMs, Mg-TAMs, lymphoid cells, oligodendroglial cells, astrocytes, neurons, endothelial cells, pericytes, and additional non-macrophage myeloid populations. To reduce over-fragmentation in the deconvolution reference, monocyte-like and monocyte-derived macrophage labels were collapsed, and lymphoid subtypes were later summed into a broader lymphocyte category for selected visualizations.

##### *6.2 Feature selection for CIBERSORTx input*

A custom 999-gene feature set was generated for CIBERSORTx mixture and reference matrices. The initial gene universe was defined as the intersection of genes present in the CED bulk count matrix, the deconvolution reference matrix, and the MSigDB Hallmark gene sets. A curated must-have gene set was then retained when available in this universe. This must-have set included canonical malignant, glial, myeloid, lymphoid, vascular, macrophage, microglial, inflammatory, and tissue-remodeling markers, including genes such as SOX2, OLIG2, PDGFRA, EGFR, GFAP, CHI3L1, PTPRC, TMEM119, P2RY12, CX3CR1, CSF1R, CD14, CD68, CD163, AIF1, PECAM1, VWF, PDGFRB, PLP1, MBP, MARCO, MSR1, IL1B, VCAN, CEBPB, ATF3, JUNB, and FOSL.

To enrich the signature matrix for genes capturing GBM cell-state programs, an additional set of genes was selected from MOFA-based factorization weights of the GBmap reference dataset, which was performed within each cell type using the mofax package with default settings as previously described (12, 13). MOFA weight matrices were filtered to retain factors with non-zero absolute loadings, and genes were ranked within each factor by signed-centered loading rank. Across cell types, mean absolute signed rank was used to identify genes recurrently represented in the latent-factor space. The union of these MOFA-selected genes and the curated marker genes was retained. The remaining candidate genes were ranked using the reference matrix by coefficient of variation after filtering for intermediate mean expression, defined as log10 mean expression between log10(10) and log10(500), and dispersion greater than 1.5. The most variable remaining genes were added until the final gene set contained 999 genes.

The CED bulk count matrix was normalized to 1,000,000 counts per sample before export as the mixture matrix. The reference matrix was subset to the same 999 genes and exported as the CIBERSORTx signature matrix. The final gene list was also written as a one-gene-per-line text file for compatibility with the CIBERSORTx interface.

*6.3 Fraction inference and post-processing*

CIBERSORTx fraction inference and expression modeling was run externally on https://cibersortx.stanford.edu/, using Hi-Resolution mode with default settings (batch correction disabled, quantile normalization disabled, and constant window size throughout deconvolution). We used the recurrent glioblastoma reference matrix as the signature matrix and the CED-topotecan bulk RNA-seq expression matrix as the mixture. The resulting sample-by-cell-type fraction table was imported back into Python. For broad cell-fraction visualizations, lymphoid subtypes were collapsed by summing T cell and Non-T lymphocyte fractions into a single Lymphocyte fraction.

Inferred cell-type fractions were visualized both as sample-level heatmaps and as group-level boxplots. For heatmap visualization, fractions were ordered by hierarchical clustering and annotated by patient and biopsy phenotype. Because inferred fractions were compositional and could vary over several orders of magnitude, visualizations used log-scaled color normalization for heatmaps where appropriate while preserving zero-valued entries as visually distinct.

Deconvolved biopsy-level cell-type fractions were analyzed using centered log-ratio transformation to account for compositionality. Fractions were normalized within each biopsy, adjusted with a pseudocount of $1\times10^{-6}$, renormalized, and transformed as $CLR(f_i)=\log(f_i)-\mathrm{mean}_j \log(f_j)$, where the mean was computed across all inferred cell types. Statistical testing was then performed on CLR-transformed fractions. For each pre-specified cell population, condition-associated differences were tested using biopsy-level linear models with patient included as a fixed blocking factor: CLR fraction ~ Condition + Patient. This model uses all biopsy-level observations while accounting for repeated biopsies from the same patient. Effect sizes represent adjusted differences on the CLR scale, with positive values indicating relative enrichment and negative values indicating relative depletion in the post-CED condition. Planned contrasts compared post-CED IN versus pre-CED, post-CED OUT versus pre-CED, and post-CED IN versus post-CED OUT. Statistical testing was restricted a priori to the cell populations emphasized in the study, including Tumor, Mo-TAM/Mo-Mono, and Mg-TAM compartments. P-values were corrected across the pre-specified cell type and contrast family using Benjamini-Hochberg FDR correction.

*6.4 CIBERSORTx high-resolution cell type-specific expression inference*

CIBERSORTx high-resolution outputs were used to approximate cell-type-specific expression profiles for selected populations in the clinical bulk cohort. High-resolution profiles were analyzed for Tumor, Mo_TAM, and Mg_TAM compartments. These inferred expression profiles were used in two ways. First, they supported heatmap visualization of cell-type-specific genes associated with infusion-zone remodeling. Second, after rescaling to approximate count space using sample-specific bulk library sizes, they were analyzed with the same PyDESeq2 framework used for the bulk count matrix as described above. Because these profiles are inferred from bulk RNA-seq, they were interpreted as supportive cell-type-resolved estimates rather than direct single-cell measurements.

**7. Murine scRNA-seq**

*7.1 Raw matrix loading and quality control*

Mouse single-cell RNA-seq data were loaded from count matrices with corresponding barcode and gene files and matched to cell-level metadata. Raw counts were stored in the counts layer before any normalization. Duplicate barcodes were removed. Quality-control metrics were computed with Scanpy after annotating mitochondrial, ribosomal, and hemoglobin genes. Cells were retained if mitochondrial transcript fraction was less than 8%, ribosomal transcript fraction was less than 8%, and hemoglobin transcript fraction was less than 5%. A minimum detected-gene threshold was defined as 100 genes, and cells below this threshold were removed. Ribosomal, heat-shock, mitochondrial, hemoglobin, and noncoding-like genes were removed from the feature matrix. Doublet detection was performed using Scrublet through scanpy.external, and predicted doublets were excluded. The filtered count matrix was normalized to 10,000 counts per cell and log1p-transformed for embedding, scoring, and visualization.

*7.2 Integrated embedding and broad cell-type annotation*

Highly variable genes were selected with a dispersion threshold minimum = 0.5 and Sample as the batch key. Principal component analysis was performed on the filtered expression matrix, followed by Harmony correction for treatment group, time point, and sample. The final Harmony-corrected PCA representation was used to construct a nearest-neighbor graph with 16 principal components, followed by UMAP embedding and Leiden clustering at resolution 2. Leiden marker genes were calculated by Wilcoxon rank-sum testing to assist manual annotation.

Neoplastic and non-neoplastic compartments were separated using tumor marker scoring, non-tumor marker scoring, and cluster-level marker gene expression. Scanpy `score_genes` was used to calculate tumor and non-tumor expression scores from curated mouse marker lists. These expression-based scores, together with differentially expressed marker genes and expected lineage markers, were used to identify malignant tumor clusters and separate tumor cells from non-neoplastic cells.

Non-tumor cells were re-embedded after Harmony correction and Leiden clustering to refine broad non-neoplastic annotations. Clusters were manually annotated using differentially expressed markers and expected lineage markers into myeloid, oligodendrocyte, astrocyte, neuron, T cell, endothelial, pericyte, and OPC-like compartments. A myeloid-restricted object was then generated for more detailed annotation of microglia-like TAMs, monocyte-derived TAMs, monocytes, and dendritic cells.

*7.3 Antunes-based myeloid label transfer*

Mouse myeloid subtypes were refined using an external myeloid reference derived from Pombo Antunes et al. GL261-derived myeloid cells (14). This dataset was used as a reference because infiltrating Mo-TAMs were distinguished from resident microglia using *CCR2*-KO. Published myeloid annotations were harmonized into the labels Mg_TAM, Mo_TAM, proliferating TAM, monocytes, and dendritic-cell subsets. A hierarchical MMoCHi classifier was trained on the reference with a hierarchy separating dendritic cells from monocyte/macrophage lineages, monocytes from macrophages, and macrophage subsets into Mo_TAM, Mg_TAM, and proliferating TAM categories (15). The trained hierarchy was applied to the project myeloid cells after intersecting the gene space between datasets.

To reduce dependence on a single stochastic classification pass, the MMoCHi label-transfer workflow was repeated across multiple independent classification runs. For each projected myeloid cell, the proportion of runs assigning each label was summarized as a membership score. Leiden clusters in the myeloid embedding were then assigned final subtype labels based on the average membership scores across their constituent cells, with proliferating TAM assignments collapsed into Mo_TAM for the final broad myeloid subtype vocabulary. This produced the final Mg_TAM, Mo_TAM, monocyte, and dendritic-cell annotations used for mouse differential abundance, MOFA, pseudotime, and pseudobulk analyses.

*7.4 Differential abundance analysis with scCODA*

Mouse cell-type differential abundance was assessed using scCODA compositional modeling. Cell counts were tabulated by sample for the final broad cell-type annotations, and each sample was annotated by treatment group and time point. Treatment was encoded with Untreated as the reference category, and time was encoded with 3-day as the reference category. The primary pan-cell-type model used the formula C(Exp, Treatment("Untreated")) * C(Time, Treatment("3d")) with Endothelial cells as the reference component. Hamiltonian Monte Carlo sampling was performed using scCODA, and credible effects were extracted across a grid of estimated FDR thresholds from 0.001 to 0.800. The same framework was repeated within the myeloid compartment using dendritic cells as the reference component to evaluate treatment- and time-associated changes among myeloid subtypes. Each cell type's FDR *p* value was reported as the minimum FDR threshold across the grid for which that cell type's effects were deemed credible by the model.

*7.5 Antunes signature scoring and Mo-to-Mg identity axis*

To further quantify myeloid-state identity, differential signatures from fine Antunes myeloid states were compiled into a mouse myeloid signature matrix. For each Antunes myeloid state, genes with the strongest positive differential expression were used to construct state-specific gene sets. These gene sets were scored in the myeloid cells with Scanpy score_genes. To generate a continuous Mo-to-Mg identity axis, Antunes-derived signature scores were standardized and projected by principal component analysis. A rotational search across two-dimensional score space was used to identify the axis that maximally separated cells annotated as Mo_TAM and Mg_TAM by standardized mean difference. The resulting axis was oriented so that Mg_TAM-enriched cells had positive values and was stored as both a raw score and a z-scored identity score for downstream visualization and pseudotime-associated analyses.

*7.6 MOFA modeling of mouse myeloid populations*

Multi-Omics Factor Analysis was used to identify unsupervised transcriptional programs within mouse Mg_TAM and Mo_TAM populations (13). For each myeloid subset, highly variable genes were selected, PCA was performed, and Harmony correction was applied for sample identity before constructing neighborhood graphs and UMAP embeddings. The subset-specific AnnData object was wrapped in a MuData object with a single RNA view, and MOFA was run with 25 factors. MOFA factor scores were extracted for cells and factor loadings were extracted for genes. Non-zero factor loadings were written as factor-by-gene weight matrices, and cell-level factor scores were written for downstream visualization, mixed modeling, and pseudotime analyses.

For Mg_TAMs, an additional analysis was performed to identify and account for resting microglia-like cells that were overrepresented in peripheral or non-tumor-containing sampled tissue. A resting signature containing genes such as P2ry12, Cx3cr1, Siglech, Tmem119, Zfp36, Nfkbia, and Csf1r and an inflamed signature containing genes such as Lgals3, Spp1, Vim, Ly6a, Tgfbi, Clec7a, and Ccl5 were scored in Mg_TAMs. A subset of resting microglia-like cells was identified using high resting-score or low-inflamed-score criteria. For the pseudotime-focused Mg_TAM analysis, variation associated with the resting/homeostatic MOFA factor was regressed out before recomputing PCA, neighborhood graphs, UMAP, and downstream pseudotime visualization. This step was used to reduce spatial sampling bias from peripheral homeostatic microglia and focus the pseudotime analysis on treatment-associated inflammatory variation.

*7.7 Diffusion pseudotime and pseudotime-associated signatures*

Diffusion pseudotime was used to order mouse myeloid cells along treatment-associated transcriptional trajectories. For the analyzed myeloid population, PCA was computed on highly variable genes, a nearest-neighbor graph was constructed from the PCA representation, and diffusion maps were calculated with Scanpy. Root cells were selected from untreated 3-day cells to anchor the trajectory in an untreated early state. Specifically, candidate root cells were restricted to Untreated_3d cells, ranked using the relevant early-state coordinate or resting-associated factor score, and the root cell was randomly selected with a fixed seed from the lowest 5% of candidate values. Diffusion pseudotime was then calculated with Scanpy DPT and z-scored across cells for visualization and downstream modeling. Enrichment of cells from each experimental condition across pseudotime was quantified by calculating a running rank-based enrichment score (ES) and comparing to randomly shuffled cells across 1000 iterations to derive empirical p-values.

To derive gene-level pseudotime-associated signatures, MOFA factor scores for cells were correlated with DPT values using Pearson correlation across cells. These factor-pseudotime correlations were then used to combine MOFA gene loadings into a single pseudotime-associated gene-weight vector. Genes with positive weights represented programs increasing along the pseudotime trajectory, whereas genes with negative weights represented programs decreasing along that trajectory. This pseudotime-derived gene vector was used as the ranking statistic for preranked Hallmark GSEA. Cell-level pseudotime-related scores, selected MOFA factor scores, the Mo-to-Mg identity axis, and Hallmark module scores summarizing proliferation, DNA-damage response, and mesenchymal/stress programs were saved for figure visualization.

*7.8 Mouse pseudobulk differential expression and GSEA*

Mouse pseudobulk count matrices were generated by summing raw counts within each biological sample and final cell type. Pseudobulk profiles were retained only when the sample-cell-type combination contained sufficient cells for stable aggregation. Differential expression was performed separately within Tumor, Mo_TAM, and Mg_TAM compartments and within each time point. Genes with fewer than 50 total pseudobulk counts across the analyzed samples were excluded. PyDESeq2 was run with treatment group as the design factor, and Topotecan was contrasted against Untreated within each cell type and time point. The resulting log2 fold-change signatures were analyzed by preranked GSEA using the mouse Hallmark library and the same permutation, seed, and size parameters described above.

#### 8. scRNA-seq from acute ex vivo slice culture

*8.1 Data loading, quality control, and integrated embedding*

The slice-culture scRNA-seq dataset comprised paired DMSO- and topotecan-treated samples from five patient-derived GBM slice cultures, including three newly generated pairs and two previously published pairs generated using the same experimental framework and reanalyzed here (16). Patient-derived GBM slice-culture single-cell RNA-seq data were loaded from a .loom file containing cell-level metadata, treatment labels, and gene-expression counts. Cells were restricted to samples from patients represented in the topotecan and corresponding DMSO controls. The raw count matrix was stored in the counts layer before normalization. Duplicate cell barcodes were removed. Quality-control metrics were computed after annotating mitochondrial, ribosomal, and hemoglobin genes. Cells were retained if mitochondrial transcript fraction was less than 10%, ribosomal transcript fraction was less than 30%, and hemoglobin transcript fraction was less than 2.5%. A minimum detected-gene threshold was determined from the lower tail of the n_genes_by_counts distribution, and cells below this threshold were removed. Ribosomal, heat-shock, mitochondrial, hemoglobin, and noncoding-like genes were removed from the feature matrix. Scrublet was used to identify and remove predicted doublets.

After filtering, slice-culture counts were normalized to 10,000 counts per cell and log1p-transformed. Highly variable genes were selected using TissueID as the batch key. PCA was performed, followed by sequential Harmony correction for Treatment and TissueID. The final Harmony-corrected representation was used to construct a nearest-neighbor graph with 16 principal components, followed by UMAP embedding and Leiden clustering at resolution 2. Cluster marker genes were identified by Wilcoxon rank-sum testing and used alongside external label transfer and marker-based classification to produce final annotations.

*8.2 GBmap label transfer and marker-based MMoCHi classification*

Slice-culture cells were annotated using both external reference transfer and marker-driven classification. For GBmap-based label transfer, a published GBmap AnnData object was loaded, raw counts were restored, duplicate gene symbols were removed. A hierarchical MMoCHi classifier was trained on the GBmap reference using a lineage hierarchy that separated tumor from non-tumor cells, immune from non-immune cells, myeloid from lymphoid cells, macrophage from non-macrophage myeloid cells, Mo_TAM from Mg_TAM cells, T cells from non-T lymphocytes, and stromal, glial, neuronal, endothelial, pericyte, oligodendrocyte, and astrocyte compartments. The trained hierarchy was then applied to the slice-culture cells in the intersected gene space without retraining.

In parallel, a marker-based MMoCHi hierarchy was applied directly to the slice-culture object. This hierarchy used canonical malignant, immune, myeloid, lymphoid, glial, neuronal, vascular, and stromal marker genes to classify cells into tumor and non-tumor lineages and then into finer lineages. Marker thresholds were run in a TissueID-aware manner, and classification was performed with the gene-expression feature limit enabled. This provided an annotation source independent of GBmap label transfer.

*8.3 CNV-aware consensus annotation*

To distinguish malignant from non-malignant cells in the slice-culture data, infercnvpy was used to calculate copy-number-like expression profiles. Human gene chromosomal positions were retrieved from Ensembl BioMart and restricted to chromosomes 1 through 22, X, and Y. Preliminary normal reference clusters were selected from confidently non-neoplastic myeloid and oligodendroglial clusters. infercnvpy was then run using these reference cells, and window-level CNV estimates were collapsed to chromosome-level summaries. For each cell, CNV metrics included correlation to the sample-specific non-normal CNV profile, global CNV signal, and a chromosome 7 minus chromosome 10 score. Tumor and non-tumor marker scores were calculated with Scanpy score_genes, and these scores were summarized at the Leiden-cluster level to support tumor-versus-non-tumor assignment.

Final slice-culture annotations were generated by consensus across three partially independent sources: Leiden/manual annotation informed by CNV and markers, GBmap-derived MMoCHi label transfer, and marker-based MMoCHi classification. Cells with full agreement were assigned directly. Remaining cells were evaluated at the broad lineage level, with rules designed to preserve malignant calls when supported by Leiden and at least one other source, to avoid forcing tumor labels when Leiden indicated a coherent non-tumor cluster and both other annotations were ambiguous, and to use agreement between GBmap and MMoCHi when present. Persistently unresolved cells were assigned by modal source label or the majority label of their Leiden cluster. Fine annotations included Tumor, Mg_TAM, Mo_TAM, Non-macrophage myeloid, T cell, non-T lymphocyte, oligodendrocyte, astrocyte, endothelial, pericyte, stromal, and neuronal labels.

*8.4 Neftel tumor-state scoring in slice cultures*

Tumor cells from the slice-culture object were scored for Neftel-like GBM cellular states using signature gene sets (10). MES-like, NPC-like, OPC-like, and AC-like scores were calculated with Scanpy score_genes. Composite MES and NPC scores were computed by averaging the corresponding paired Neftel signatures (10). For visualization of acute tumor-state changes after topotecan treatment, signature scores were averaged by TissueID and treatment condition and compared between matched DMSO and topotecan-treated slices. For discrete tumor-subtype assignment, standardized Neftel state scores were transformed into a two-axis decision space separating OPC/NPC-like from MES/AC-like states and then assigning cells to NPC-like, OPC-like, MES-like, or AC-like categories according to the sign and magnitude of the corresponding state contrasts. Groups were compared using a paired Student's t test with $p < 0.05$ considered significant.

*8.5 Slice-culture differential abundance analysis*

Differential abundance of slice-culture cell types was analyzed with scCODA (17). Cell counts were tabulated for each TissueID-treatment pair and final fine annotation. Each sample was annotated by treatment, and treatment was modeled with DMSO as the reference category. The scCODA model used Treatment as the design formula and Stromal cells as the reference component. Hamiltonian Monte Carlo sampling was performed, and credible effects were extracted across a grid of estimated FDR thresholds from 0.001 to 0.800. For each cell type and treatment coefficient, the effect estimate, posterior standard deviation, credible-effect status, and minimum FDR threshold at which the effect was detected were summarized for plotting.

*8.6 Slice-culture pseudobulk differential expression*

Slice-culture pseudobulk profiles were generated by summing raw counts within each TissueID, treatment condition, and final cell type. For tumor analyses, counts from MES-like, AC-like, NPC-like, and OPC-like tumor states were summed to create a broad Tumor pseudobulk profile for each TissueID-treatment pair. For myeloid analyses, Mg_TAM and Mo_TAM pseudobulk profiles were analyzed separately. Rows with zero total counts were excluded, and genes with fewer than 100 total counts across the analyzed pseudobulk samples were removed. For each retained cell type, only TissueIDs represented in both DMSO and the drug condition under analysis were included. PyDESeq2 was run with TissueID and Treatment as design factors, and the drug condition was contrasted against DMSO. Slice-culture pseudobulk differential-expression signatures were analyzed by preranked GSEA using log2 fold change as the ranking statistic. Hallmark and custom-signature enrichments were calculated for Tumor, Mg_TAM, and Mo_TAM topotecan signatures.

**9. Bulk RNA-seq analysis of iPSC-derived microglia treated with topotecan or LPS**

Bulk RNA-seq analysis of induced pluripotent stem cell-derived microglia was performed to compare topotecan-induced transcriptional remodeling with a canonical inflammatory stimulus. Bulk RNA-seq differential-expression results for iPSC-derived microglia were imported from DESeq2-derived output tables. Differential-expression tables were parsed for topotecan-treated and LPS-treated iPSC-derived microglia relative to control (DMSO), and log2 fold changes and adjusted p-values were combined across contrasts. For each condition, a differential-expression table containing gene-level log2 fold change and adjusted p-value was exported as source data.

Volcano plots were generated from these differential-expression results by plotting log2 fold change against -log10 adjusted p-value. Genes exceeding the specified fold-change and significance thresholds were colored according to the direction of change, and the most significant or largest-magnitude genes in each direction were labeled. Hallmark and custom-signature GSEA were performed on the iPSC-microglia topotecan and LPS signatures using the same preranked GSEA framework applied to the clinical, mouse, and slice-culture signatures.

#### SECTION C – STATISTICS

##### 10. Statistics and reproducibility

*10.1 Statistical conventions*

Bulk RNA-seq and pseudobulk single-cell differential expression were analyzed using negative-binomial models in PyDESeq2. Single-cell differential abundance was analyzed using scCODA compositional models. Quantitative immunofluorescence and biopsy-level summaries were analyzed using mixed-effects or robust regression models where repeated measurements were present, as described in the relevant sections. P-values were adjusted for multiple testing using Benjamini-Hochberg FDR correction, Holm correction, or Bonferroni correction depending on the analysis.

*10.2 Differential expression and enrichment statistics*

Differential expression analyses were performed at the sample or pseudobulk level. Bulk human biopsy comparisons modeled raw counts with patient identity and biopsy phenotype. Slice-culture pseudobulk comparisons modeled raw counts with patient/tissue identity and treatment condition. Mouse pseudobulk comparisons modeled raw counts within each cell type and time point using treatment condition. For all DESeq2-style analyses, adjusted p-values were calculated with the Benjamini–Hochberg method. Gene set enrichment analyses were performed with preranked GSEA on log2 fold-change vectors, and pathway-level false discovery rates were used to determine significant enrichment. Cell-type abundance analyses used scCODA for compositional modeling in single-cell datasets, supplemented by biopsy-level and patient-aware fraction analyses for CIBERSORTx-derived bulk deconvolution estimates.

*10.3 Software environment*

All analyses were performed in Python using pandas, numpy, scipy, scanpy, anndata, scanpy.external, muon, mofax, statsmodels, seaborn, matplotlib, mmochi, infercnvpy and related scientific Python dependencies. Analyses were performed across two Python environments. The primary computational environment used for source data processing, latent-factor modeling, and external projection workflows was hosted on an Amazon EC2 instance running Amazon Linux 2023 with Python 3.10.19 in a micromamba environment. Key packages in this environment included scanpy 1.11.5, anndata 0.11.4, muon 0.1.7, mofapy2 0.7.3, harmonypy 0.2.0, infercnvpy 0.6.1, liana 1.7.1, scvi-tools 1.3.3, ctxcore 0.2.0, pandas 2.3.3, numpy 1.26.4, scipy 1.15.2, statsmodels 0.14.6, scikit-learn 1.7.2, matplotlib 3.10.8, and seaborn 0.13.2. Additional local analyses, including annotation-related workflows and downstream exploratory analyses, were performed on a macOS arm64 system using a conda environment with Python 3.8.18. Key packages in this local environment included scanpy 1.9.6, anndata 0.9.2, muon 0.1.6, mofapy2 0.7.1, mmochi 0.2.2, pandas 1.5.3, numpy 1.24.4, scipy 1.10.1, statsmodels 0.14.0, scikit-learn 1.3.2, matplotlib 3.7.3, and seaborn 0.12.2.
